# Cerebellar connectivity maps embody individual adaptive behavior

**DOI:** 10.1101/2021.02.24.432563

**Authors:** Ludovic Spaeth, Jyotika Bahuguna, Theo Gagneux, Kevin Dorgans, Izumi Sugihara, Bernard Poulain, Demian Battaglia, Philippe Isope

## Abstract

From planification to execution, cerebellar microcircuits encode different features of skilled movements. However, it is unknown whether cerebellar synaptic connectivity maps encode movement features in a motor context specific manner. Here we investigated the spatial organization of excitatory synaptic connectivity in mice cerebellar cortex in different locomotor contexts: during development and in normal, trained or altered locomotor conditions. We combined optical, electrophysiological and graph modelling approaches to describe synaptic connectivity between granule cells (GCs) and Purkinje cells (PCs). Synaptic map maturation during development revealed a critical period in juvenile animals before the establishment of a stereotyped functional organization in adults. However, different locomotor conditions lead to specific GC-PC connectivity maps in PCs. Ultimately, we demonstrated that the variability in connectivity maps directly accounts for individual specific behavioral features of mice locomotion, suggesting that GC-PC networks encode a general motor context as well as individual specific internal models underlying motor adaptation.

## Introduction

Sensorimotor adaptation and motor learning rely on a combination of neuronal computation performed in different brain areas embedded in multiple cerebello-cortical loops^1–5^. The population dynamic of cortical networks encodes stimulus features, which are communicated to different topological nodes of the brain dedicated to motor planning and motor control. If these neuronal dynamics succeed in producing an adapted behavior, they are stabilized in specific areas through synaptic plasticity yielding specific routing of action potentials across the network^6–9^. Recent experiments described how different types of neurons in the cerebral cortex are selectively connected depending on stimulus features they have to encode^10–13^. Similarly, in the cerebellar cortex, stereotyped albeit plastic and structured connectivity maps across individuals have been described in identified modules^14^. However, what connectivity maps encode precisely in the cerebellar cortex is still unclear. In a first scenario, connectivity maps reflect features (e.g., limb movement or velocity) that could be used for context specific sensorimotor adaptation by other brain areas (e.g., during walking, running, or swimming). Maps would be context-independent as an invariant internal model of the features^15, 16^. In this scenario, plasticity would be essentially related to modification of relative body coordinates (e.g., limb size during development) and maps should not be modified in different contexts. Alternatively, in a second scenario, a given map would encode features in a specific context (e.g., limb movement when running). Connectivity maps would represent individual-specific engrams of adaptive behaviors established throughout motor learning and such feature-based maps should be animal and context specific. Experiments suggesting that the cerebellum stores in its neuronal network internal models of the body predicting an expected sensory feedback of motor commands are compatible with both scenario ^17–22^.

We therefore investigated whether cerebellar connectivity maps are modified during development and by altering locomotor contexts. We targeted the anterior vermal lobule III-IV, an area involved in postural adaptation of locomotion^23–25^ as pointed out by gait impairment and altered locomotion occurring after lesions or alteration of cerebellar plasticities in the cerebellum^23, 26, 27^. Functional studies have identified task-related modules involved in specific movements and described the cerebellum as an array of anatomo-functional modules and microzones defined by the topographical organization of a major excitatory input of the cerebellum, the climbing fibers (CF) ^28–31^. Microzones are composed of a group of PCs, the sole output of the cortex, having similar CF receptive fields and can be identified through differential expression of molecular markers (known as Zebrins) in PCs^30, 32–35^ (Fig 1a). PCs combine information from excitatory granule cells (GC) relaying mossy fiber inputs (MFs) and from the CF which control plasticity at the GC-PC synapses ^36–39^. Sensorimotor MF inputs are multimodal and project to many different patch of GCs in different lobules, defining a fractured somatotopy ^40–42^. Furthermore, GCs that receive MFs from different modalities^43, 44^ compute and distribute this information to many different microzones via their long axons, the parallel fibers (PFs) leading to specific excitatory connectivity maps patterns conserved between animals^14^. Therefore, GC-driven synaptic connectivity maps define how cerebellar modules are coordinated in a given cerebellar lobule and may encode common and generic adapted behavior ^14, 41^.

**Figure 1:**
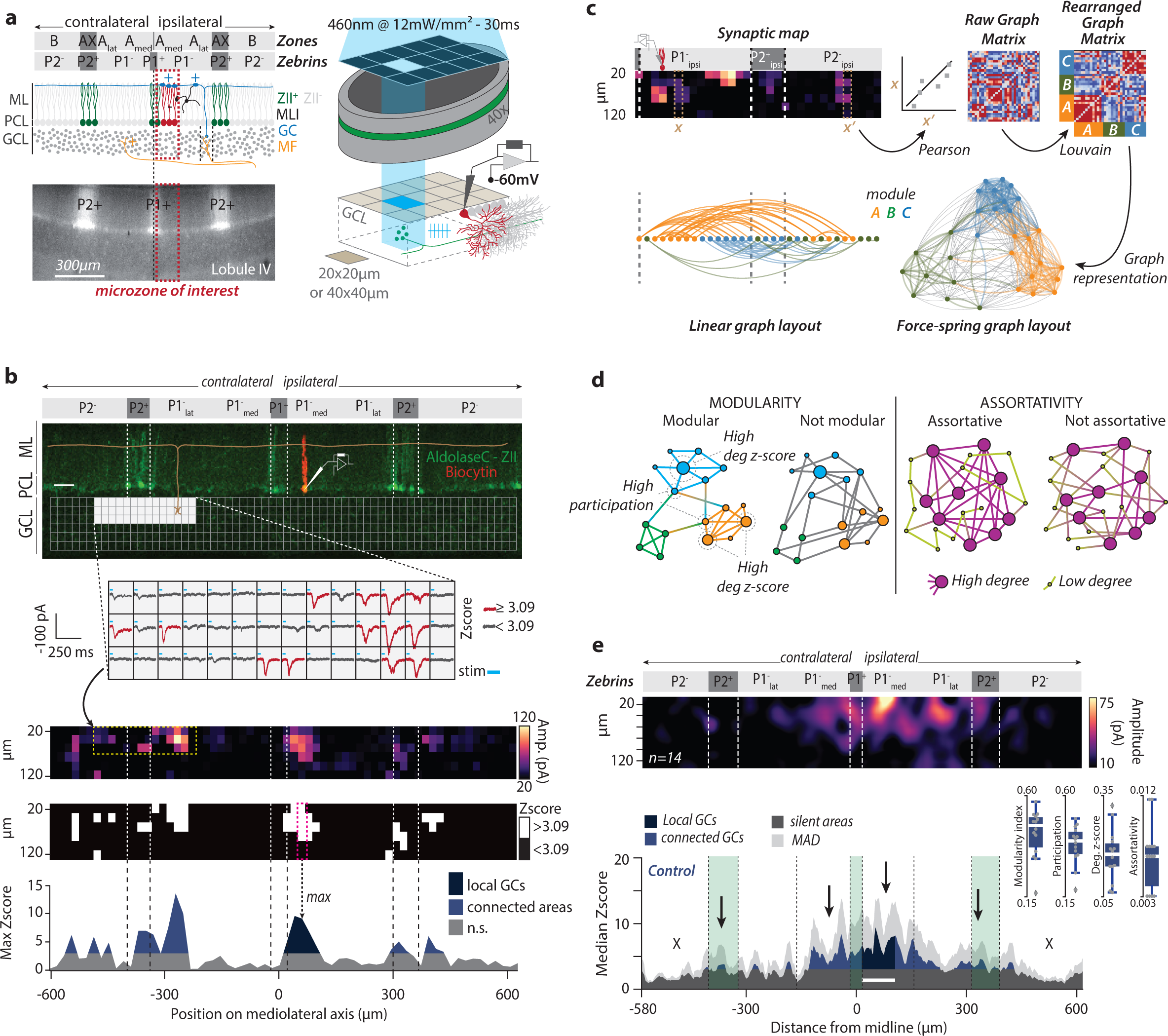
Stereotyped and structured GC-PC connectivity maps. **(a) *Upper left panel***, diagram illustrating cerebellar cortical anatomy in brain slices. Recorded PCs (*green and red*) are close to the midline (>120 μm) and belong to the P1-medial microzone in the A zone of lobule III/V. ZebrinII positive (ZII+) and ZebrinII negative (ZII-) PCs coincide with some microzonal boundaries. GCs relay MF multimodal information to PCs across cerebellar microzones via monosynaptic excitation or disynaptic inhibition through molecular layer interneurons (MLIs). ***Bottom left panel***, picture of fluorescent PCs in an acute cerebellar slice from an ALDOC-Venus mouse. ***Right panel***, photostimulation apparatus for glutamate uncaging. Systematic release of caged-glutamate was done in 20x20 or 40x40μm GCL areas using patterned-light illumination while PCs are recorded in voltage-clamp (holding=-60mV). *GCL, granule cell layer; ML, molecular Layer; PCL, Purkinje cell layer*. **(b)** Example of GC-PC synaptic map. ***Upper part***, post-hoc reconstruction of a recorded PC. *Scale bar = 50μm*. Light-evoked EPSCs from each stimulation site are averaged. ***Lower part***, resulting synaptic maps are translated in z-score maps (methods and Fig.S2). The maximal z-score value for each GCL column (*pink-dashed box*) along the mediolateral axis is projected on a 1D map to define GC synaptic patterns. Blue areas show GC areas connected to the recorded PC (z≥3). **(c)** Connectivity map preprocessing for graph network analyses. Column-wise Pearson correlations (e.g., *x and x’ columns*) of synaptic weights lead to a spatial correlation matrix (“*Raw graph matrix*”), highlighting GCL columns eliciting similar responses in the recorded PCs. The raw matrix was rearranged using the Louvain community detection algorithm giving the “*Rearranged graph matrix*” with well-defined graph modules (A, B, C). Modules are visualized using a force-spring algorithm in an arbitrary space (*Force spring layout*, i.e., distance between nodes represents the degree of correlation and the colors represent the module identity from the “*Rearranged matrix*”) or according to nodes positions along the mediolateral axis (*linear layout*). **(d)** Graph parameters. The module degree z-score illustrates how a given node is connected to nodes from the same module while participation illustrates how uniformly a node is connected to nodes from different modules. Assortativity illustrate the potency of nodes to connect with nodes having similar degree. **(e)** GC-PC connectivity maps of the CTRL group of mice (n=14, see Table 1). ***Upper panel***, averaged 2D map. ***Lower panel***, median 1D pattern. White bar, averaged PC position. Inset, distribution of graph parameters measured in CTRL graphs.

We, here, use a combination of electrophysiological recordings and systematic glutamate uncaging to establish GC-PC connectivity maps in different locomotor contexts and development. We then describe map spatial organization using a novel graph-based modelling of synaptic weights. Through functional reconstruction, graph representations and the quantification of their graph-theoretical features, we showed that the behavioral context leaves a trace in the spatial organization of maps. We also found that while connectivity maps are correlated to behavioral conditions, each mouse developed a specific individual combination of connectivity traits linked to its individual and highly specific locomotor activity, suggesting that behavior may causally shape the spatial (re)organization of connectivity maps.

## Results

### Stereotyped synaptic transmission in the GC-PC pathway

We established synaptic connectivity maps between GCs and PCs in lobule III-IV of the anterior vermis and selected a group of medial PCs (0-120 µm, medial P1- band of the A zone^14^; Fig.1a, Fig.S1) which are involved in postural adaptation during locomotion. PCs from this microzone were whole-cell patch-clamped on acute cerebellar slices from AldolaseC- Venus fluorescent transgenic mice^45^ (Fig.1a; N=83 mice, Table 1) and inhibition was blocked. *In situ* fluorescence of AldolaseC-venus (Zebrin II) in PCs allowed direct visualization of cerebellar microzones and accurate targeting of the medial PCs (*microzone of interest* in Fig 1a). Rubi-Glutamate^46^ was sequentially photostimulated on the granule cell layer (GCL) following a grid of 128 or 384 square sites (20x20 or 40x40µm corresponding to high vs low resolution, respectively) while EPSCs were recorded in PCs (n=148 PCs, Fig.1a,b, S1, S2 and Table 1; see Methods). For each PC, a single GC-PC connectivity map was built from the averaged EPSCs elicited by each of the 128 or 384 photostimulated GCL sites (Fig.1b, S2, each average built from 6-10 mappings). To define whether a site is functional or silent, EPSC amplitudes were normalized to noise level and expressed as z-scores (Fig.1b, S2 and methods). MF inputs originating in a specific precerebellar nuclei^14, 42, 47, 48^ project onto GCs at several location in a lobule and in most of the GCL height, defining a fractured and patchy columnar somatotopy (Fig S1; see^14, 40, 41^). We therefore treated the GCL either as a grid of 384 or 128 GC sites or as a collection of GC columns of 20 or 40-µm width, respectively. Each connectivity map was then represented either as a 2D map or a projected 1D pattern corresponding to the maximal synaptic weight of GC columns (Fig.1b and Methods).

**Table 1:** Group description.

| Condition | Acronym | N<br>(mice) | n<br>(maps) | Description |
| --- | --- | --- | --- | --- |
| Pups | PND9-10 | 5 | 11 | Mice raised in standard conditions |
| Juvenile | PND12-13 | 6 | 12 |  |
| Adolescent | PND14-18 | 7 | 15 |  |
| Adult | PND>30 | 7 | 10 |  |
| Control | CTRL | 9 | 14 | Mice raised in standard conditions |
| Short Training | STR | 6 | 12 | 7 daily sessions in the running wheel |
| Long Training | LTR | 6 | 11 | 10 daily sessions in the running wheel |
| Early Cuff | EC | 13 | 25 | Cuff implant on the right sciatic nerve |
| Early Sham | ES | 7 | 10 | Cuff surgery only |
| Late Cuff | LC | 8 | 16 | Cuff implant on the right sciatic nerve |
| Late Sham | LS | 9 | 12 | Cuff surgery only |
| **Total** |  | **83** | **148** |  |

We also developed an analytical method allowing us to investigate the “patchiness” of GC-PC connectivity maps that could consider the fractured MF projections projecting onto distant connected GC patches. To do this, we used an analytic workflow based on mathematical graph representation of the connectivity maps (Fig.1c, d, methods). Graphs (or networks) are generic abstract entities composed of nodes and links between them^49^. We considered each connectivity map as a matrix *M*, whose *M_xy_* entry denotes the EPSC amplitude recorded at the GC site coordinates (x,y) (Fig.1c). We computed the normalized Pearson correlations (*C_xx_’*) for every pair of columns (mapped to graph *node*s) *x* and *x’* along the mediolateral axis of the connectivity maps. *C_xx_’* entries become the strengths (i.e., weights) of links between graph nodes associated to the positions *x* and *x’. Cxx’* values were compiled in a correlation matrix *C(M)* (*“Raw Graph matrix”* on Fig.1c) which can be considered as the adjacency matrix of an undirected graph (Methods). Map columns traversed by one or multiple patches will have similar synaptic profiles, and the corresponding graph nodes will thus be strongly connected, forming *network modules*. These modules are highlighted as matrix blocks, appearing in the adjacency matrix after sorting its rows and columns according to a Louvain community detection algorithm^50, 51^ (“*Re-arranged Graph matrix*” in Fig.1c). Hence, spatially distant GC sites can belong to the same graph module. In Fig.1c, nodes belonging to 3 different modules have been colored into two alternative visualization of the graph associated to a map. A first linear layout, in which node positions follow the original mediolateral axis alignment, manifests that modules tend being made of spatially contiguous nodes, with some exceptions (spatial distance between nodes in a same module is ∼10% smaller on average than distances between modules). We show as well –and prefer in the following– an alternative network layout, optimized for module visualization by a force-spring algorithm, which reduces (increases) the distance between the strongly (weakly) connected nodes.

We quantified the structuration of graph networks using 4 standard metrics (Fig.1d, see methods). (1) The *modularity index* is a measure of how modular or “patchy” is a graph. (2) The average *module degree z-sco*re which can be used as a marker of degree heterogeneity within modules. (3) The average *participation coefficient* measures the probability that a node in a module is also connected to nodes in other modules. (4) The average local *assortativity* measures the tendency of nodes to connect to other nodes with similar strength (i.e., the sum of weights of connections to a node). Together these abstract metrics convey concrete and complementary information about the geometry of the connectivity maps: the larger the modularity, the larger the general degree of “patchiness”; the larger the degree z-score, the more intricate the patch shapes; the larger the participation, the more overlapping the patches; the larger the assortativity, the clearer the separation between small and big patches. Using connectivity maps and graph properties, we assessed whether and how the maps are modified at different stages of postnatal development (pups, juvenile and young adults), after an injury (right-hindlimb impairment via sciatic nerve cuffing) and after locomotor training in a running wheel (see methods and Table 1).

First, we established high resolution connectivity maps in naive adult animals (CTRL, n=14 maps, N=9 animals, PND>30, Table 1, Fig.1e). Medial PCs individual maps were composed of GC sites eliciting a wide range of responses from silent sites to strong connections resulting in EPSCs of hundreds of pA (mean±SD = 72.3±49.8 pA, from 0 to 393.2 pA). Local and distant patches of connected GCs interleaved with silent GC areas were systematically observed (Fig.1e) and the distribution of connected GC sites was skewed in the GCL depth (Fig.S3a). Building the median of all the 1D patterns of connectivity (Fig.1e, methods) we showed that in CTRL condition at least 50% of the patterns lacked functional GC sites in ipsi- and contralateral P2^-^ and P1^-^_lateral_ bands while most of the connected GC sites were observed in P1^-^_medial_ and P2^+^ bands (ipsi/contralateral). To assess the robustness of the median 1D pattern, we used a bootstrap analysis after shuffling 1D patterns (Fig.S4). This analysis confirmed that connectivity maps are not randomly distributed but rather shared a similar GC-PC synaptic organization across animals as shown in our previous study^14^. Hence, the stereotyped patchy organization of GC-PC excitatory connectivity maps suggests specific microzonal communication. The existence of a non-random organization with patches simple in shape, imperfectly separated and with a continuum spectrum of possible sizes is suggested as well by graph representations, which display a relatively large modularity (0.43±0.1, inset in Fig. 1e) and average participation coefficient (0.38+0.09), while averaged module degree z-scores (0.17+0.08) and assortativity (0.007±0.003) are low.

### Network connectivity rules mature during development

We first addressed how GC-PC connectivity maps were established and structured during postnatal development (i.e., in a normal adaptive locomotor context). Since the motor apparatus and motor coordination undergo progressive adaptation, especially when animals open their eyes and start walking^52–54^, we assessed whether conserved connectivity maps observed in adults originate from a linear and progressive functional wiring. If connectivity maps reflect the strict establishment of the input/output relationships in a topographic manner, we expect to observe a corresponding structuration of the patchy microzonal organization. We then established GC-PC connectivity maps at different time point during mouse development: from PND9-10 before proper quadrupedal locomotor activity (mice crawl, eyes are closed) to young adults (PND>30) when locomotion is well adapted (Fig.2, Table 1).

**Figure 2.**
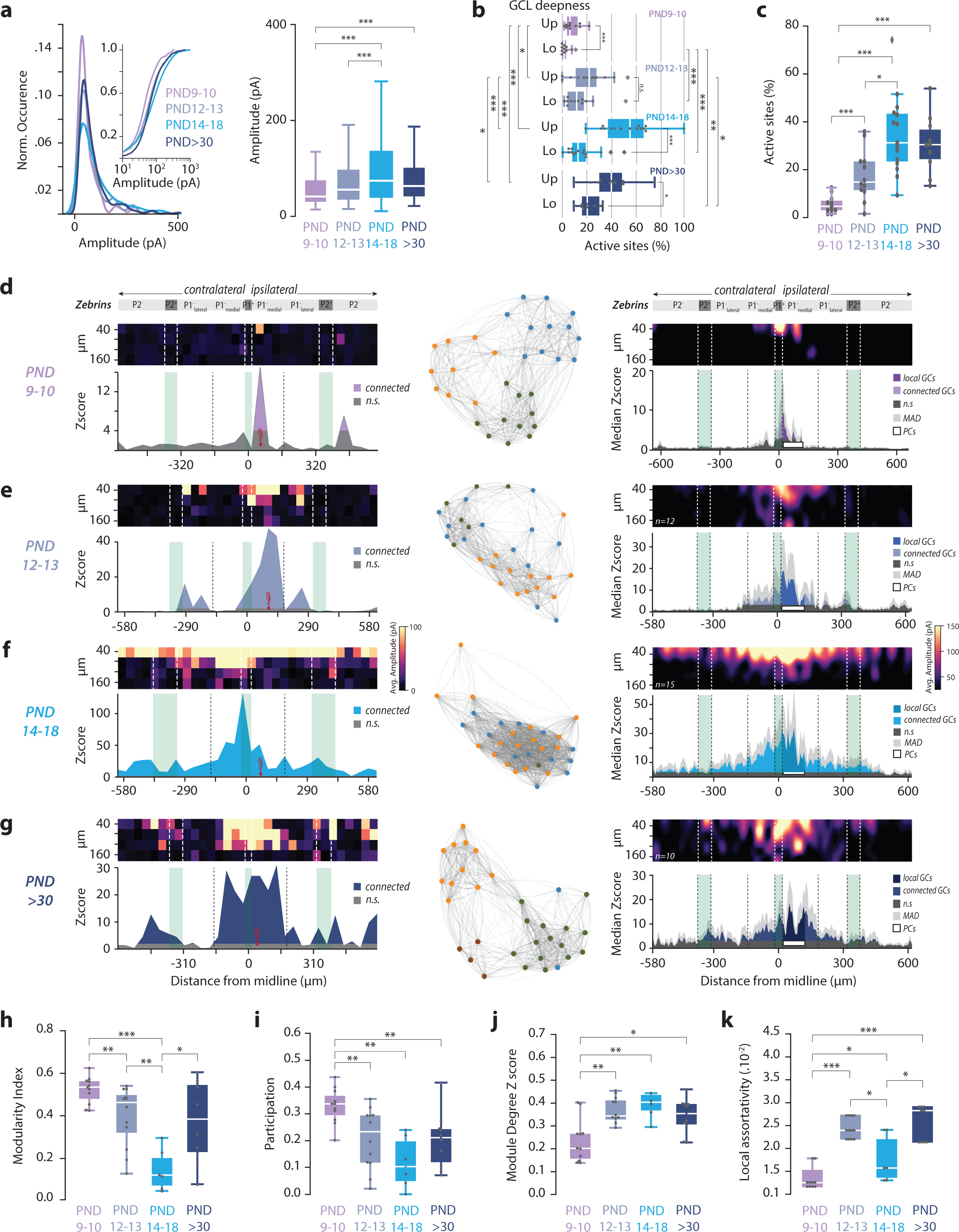
Postnatal development of connectivity maps. **(a)** Distribution of GC-PC significant synaptic weights during postnatal development (PND 9-10, 12-13, 14-18, 30-40); ***left***, normalized and cumulative representation (*inset*); ***right,*** boxplots. For statistics, see table in Figure S10. **(b)** Proportion of GC active sites in GCL height. (*Up*), upper GCL, 0-80µm; (*Lo*), lower GCL, 80-160µm. For statistics, see table in Figure S10. **(c)** Proportion of GC active sites measured in the total GCL of GC-PC maps. For statistics, see table in Figure S10. **(d, e, f, g)** Connectivity maps at PND9-10, PND12-13, PND14-18 and adult mice. ***Left column***, examples of GC-PC map; ***middle column*** corresponding graph plots; ***right column***, average 2D maps and median 1D patterns. Median absolute deviation (MAD) is shown in light grey. **(h, i, j, k)** Graph properties (modularity, participation, module degree z-score and local assortativity) of GC-PC synaptic maps at PND9-10, PND12-13, PND14-18 and adult mice. Average values and statistical comparison are availiable in Figure S10.

At PND9-10, low-resolution photostimulation triggered EPSCs (mean = 62.1±55.4 pA, Fig.2a) mostly from GCs located below the recorded PC (Fig.2b). Connected GC sites represent on average 5.5±3.6% of the total illuminated area (Fig.2c, d). However, distant GC inputs were already observed (Fig.2d) as PF lengths in the medial vermis already exceed several hundred µm long (Fig.S5a). Graph parameters are similar to the CTRL groups suggesting that synaptic maps are already patchy, unstructured internally and overlap between patch boundaries (modularity=0.52±0.06; participation=0.33+0.06; module degree z-scores=0.23+0.09; assortativity=0.014±0.002; Fig 2h-k).

At PND12-13 (just before eye opening ^52–54)^ we observed a three-fold increase in the number of active sites (PND12-13: 17.6±9.9% vs PND9-10: 5.5±3.6%, p=0.0148, Mann-Whitney U test, Fig.2c) and larger EPSCs (mean EPSCs PND12-13: 87.3±94.2pA vs PND9-10: 62.1±55.4pA, p=0.011, Mann-Whitney U test, Fig.2a). This increase in connectivity illustrates the appearance of new functional GC-PC synapses rather than a change in GC excitability as no difference could be observed in GC firing rates between pups and PND12-13 following glutamate uncaging (Fig.S5b). With maturation, modularity and participation decreased while module degree z-score and assortativity increased illustrating a stronger internal structuration of the modules (i.e., graph modules are less compact, better defined and overlap less, Fig.2e Fig S10).

PND14-18 epoch corresponds to a very active motor period in mice when animals open the eyes and start walking^53^. Strikingly, GC inputs were spread all along the mediolateral axis from P1^+^ to P2^-^ bands (Fig.2f) with a significant increase in the amount of active GC sites (PND14- 18=34.7±15.6% vs PND12-13=17.6±9.9% at, p=0.03008, Mann-Whitney U test, Fig.2c), particularly in the first upper part of the GCL (mean=57.7±20.5%, Fig.2b). Concurrently, modularity, participation and assortativity indexes reached a minimal value (0.15±0.08; 0.1±0.08; 0.02±0.005 respectively, Fig.2h, I, Fig S10) while the module degree z-score was maximal (0.41±0.08, Fig.2j, and S10). These results indicate that the limits between patches became blurry, with responses organized around a center with maximum amplitude of evoked response (confirmed by low modularity index at PND14-18: 0.15±0.09 when compared to PND9-10: 0.52±0.06; PND12-13: 0.4±0.13 and PND>30: 0.37±0.18, p=0.0005; 0.002 and 0.01 respectively, Mann-Whitney U test, Fig.2h). These findings suggest a loss of network structure around this critical age.

Beyond PND30 while the number of active sites remain stable (30.6±11.4% vs 34.7±15.6%, p=0.99, Mann-Whitney U test, Fig.2c), connectivity maps resumed to a more refined patchy organization (Fig.2g). Indeed, modularity increased again for most maps (0.38±0.18, Fig.2h, S10) corresponding to a re-emergence of multiple patches, as before PND14, but with stronger internal organization, as visible from the smaller average participation coefficient (0.2+0.1, Fig.2i and S10) and higher average degree z-score (0.34±0.07, Fig.2j and S10) and assortativity (0.026±0.004, Fig.2k and S10). Therefore, as suggested by connectivity maps, the analysis of graph properties showed that network properties do not mature linearly with age. Rather, they first display dense and unstructured connectivity in young PND14-18 animals, then they evolve toward the appearance of a patchy well-structured connectivity in adulthood. Altogether, these results suggest that connectivity maps are not built solely by following strict anatomical input/output rules, but likely rely on adaptive mechanisms occurring during development.

### Adapted locomotor activity correlates with specific connectivity maps

Development leads to conserved adult connectivity maps. We now asked whether different locomotor contexts yielding specific locomotor adaptations affect synaptic maps organization in adults. We performed high-resolution connectivity mappings (20x20µm/GCL site) in mice that underwent different locomotor adaptive behaviors (see Table 1): (1) two groups of mice were trained (TR) to run on a wheel (1h/day) for 7 (STR) or 19 (LTR) days (Fig.3a) learning a new type of gait, gallop^55^. In the last session, LTR mice have traveled 2.8-fold more distance compared to first session (mean distance D1: 331±107m; D19: 932±97m, paired t-test between D1 and D19, p=2.10^-4^; Fig.3a). Connectivity maps were established at 7 or 19 days of training time. (2) In other groups, locomotion was impaired by inserting a cuff around the sciatic nerve of the right hindlimb (Fig.3b) and connectivity maps were established at 2-9 days (early cuff, EC) or after 21 days (late cuff, LC) after the surgery to allow animals to fully re-adapt. Two sham groups (surgery only) were also included (*early sham*, ES and late sham, LS).

We evaluated locomotor deficits in cuffed and sham mice on a corridor equipped with two streams of force sensors to monitor weights from either side of the body when walking (see methods). The balance index (BI) is the ratio of the total weight generated by left vs right side of the body along the corridor (Fig.3b; Methods). The BI was measured during one month after cuff or sham surgeries before connectivity maps were recorded in acute slices. As expected, before surgery BI index was close to 0 in all mice (Cuff, -0.04±0.20; Sham, -0.11±0.15; CTRL, -0.23±0.17). After surgery balance was altered in Cuff and Sham animals (p=0.0433, F=2.16, one-way RM-MANOVA, Fig.3c). In the first few days after surgery, sham and cuff mice limp on the right side and the negative BI indicate a higher weight on the side of the surgery (Cuff, - 0.45±0.09; Sham, -0.37±0.08; Ctrl, 0.12±0.13; p=4.45.10^-3^, F=7.18, one-way ANOVA at D4, “*Early phase”,* Fig.3c). The following week, while mice were not limping anymore, sham and cuff mice did not recover full balance. The BI switched to a positive value, illustrating compensatory behavioral strategies, with a peak 15 days after surgery (Cuff, 0.82±0.13; Sham, 0.27±0.12; Ctrl, -0.06±0.19; p=2.47.10^-3^, F=8.23, one-way ANOVA at D15, “*Late phase”,* Fig.3c). Ultimately, after one month all mice recovered balance and no apparent locomotor impairment remained (Cuff, 0.11±0.06; Sham, -0.01±0.05; Ctrl, 0.10±0.09; p=0.368, F=1.052, one-way ANOVA at D28, “*Recovery phase”,* Fig.3c). While we observed that sham animals (LS) quickly recovered their ability to walk after the surgery, balance recovery followed a similar time-course than cuffed animals albeit with a smaller maximal impairment (area under the curve at D15, Cuff, 1.27±0.76; Sham, 0.59±0.88; Fig.3d, p=0.0226, Mann-Whitney U test).

Connectivity maps were established in acute slices either before locomotor adaptation (ES and EC) or after full recovery (LS, LC, or TR). We first analyzed the overall distribution of the synaptic weights (Fig.4a). Synaptic weights in active sites (z-score>3.09) were significantly larger in LC, Sham and TR animals compared to CTRL group (Kruskal-Wallis test, p=1.10^-71^; Fig.4a and S10). Furthermore, the mean synaptic weights increased with the time post-surgery in cuffed and Sham animals (EC vs LC, p=5.10^-12^; ES vs LS, p=2.10^-15^; Mann-Whitney U test, Fig.4a and S10). While the proportion of active sites per full map were not significantly different between conditions (Fig.4a and S10, p=0.132159, Kruskal-Wallis test) we found regional differences when comparing from local vs distal GCL inputs in all conditions but CTRL (Fig.4b, S3b, c, d). Similarly, synaptic amplitudes were larger in local vs distal GCL areas in EC, LC, and TR animals, but not in sham (Fig.4b and S10, S3a). Altogether, these findings indicate that long-term locomotor adaptation either promotes the appearance of new active synapses or potentiates existing ones in connected GC sites at specific locations.

We then analyzed the spatial organization of GC connectivity to medial PCs across all conditions. As in the CTRL condition (Fig.1e), all the averaged connectivity maps established using photostimulation showed a clear patchy organization (Fig.4c). Median synaptic patterns showed that local GCs systematically elicited significant inputs and showed distal hotspots and non-connected areas at the exception of LC animals in which connected GC sites were observed in most GCL columns. Each condition yielded a specific conserved median pattern different from the CTRL group (STR, p=7.10^-7^; LTR, p=2.10^-3^; ES, p=8.10^-10^; LS, p=8.10^-06^; EC, p=4.10^-08^; LC, p=7.10^-18^, Kolmogorov-Smirnov test). While connected GC sites were rare in ipsi and contralateral P2^-^ bands in CTRL and LS animals, 1 or 2 hotspots were observed at a similar location in ipsilateral P2^-^ band (typically 450-600µm from the midline) in TR (S and LTR) and cuffed (EC and LC) animals. Connectivity also differed in the P1^-^_lateral_ and P2^+^ bands: in CTRL, few GC sites were connected in P1^-^_lateral_ bands; in TR animals, connected sites appeared in STR and strengthened in LTR in contralateral P1^-^ band. In LS, both ipsi and contralateral P1^-^ bands were also highly connected to medial PCs. Conversely, while P2^+^ bands were both connected in CTRL, connections disappeared in ipsilateral P2^+^ bands in L- TR and in contralateral P2+ bands in LS. In addition, 2D maps revealed an increased connectivity in the upper part of the GCL, especially in LC mice (Fig.S3a). Altogether, these results demonstrate that different adapted contexts of locomotion directly influence connectivity maps in the cerebellum. However, in a given locomotor context, maps do share a common architecture across animals, suggesting that the synaptic re-organization might operate by selecting new condition-specific MF inputs relaying information from different muscles or limbs.

### Graph descriptions of connectivity maps but not synaptic weights can discriminate different locomotor adaptation conditions

The median distribution of the 1D patterns illustrates the overall probability of connection between medial PCs and distant microzones, however, the distribution of the median absolute deviations (MAD; Fig.4c) suggests that despite common architecture, non-negligeable variability remains between animals in the same group. To quantify this variability (intra-condition) we assessed whether synaptic weights in identified microzones co-varied between slices and animals. Synaptic weights from all active GC sites in each microzone were averaged and pairwise linear regressions between microzones were performed across animals in each condition (see methods). We found specific pairs of correlated microzones in each locomotor condition (Fig.5a). Notably, we observed a 5-fold increase in the number of correlated bands in the LC condition, the most invasive protocol (Fig.5b, S6a). These findings suggest that the relationships between connected microzones and medial PCs are conserved between slices and animals, which may illustrate common integration of specific MF inputs.

To evaluate the robustness of the condition-specificity of the connectivity maps we tested whether single maps could be classified in their underlying locomotor condition by considering only microzonal synaptic weight distribution. Connectivity maps were first vectorized in an 8- dimensional space, each dimension being the average synaptic weight found in a microzone (8 microzones). We used a non-linear dimensionality reduction algorithm (t-Stochastic Neighbor Embedding technique, t-SNE, Methods, Fig.5c) to possibly visualize clusters of maps with similar synaptic weight profiles in our dataset. When maps were labelled by conditions (L/EC, L/ES, TR and CT), 2D projections showed strong scattering and overlapping center of gravity (group medians of tSNE projection coordinates, Figure 5c), indicating a poor discrimination of behavioral groups. We also used a supervised machine-learning algorithm (random forest classifier^56^, methods) to predict one of the possible adaptive condition labels based on the same vectors of average synaptic weights used as inputs. A confusion matrix in which rows represent the actual condition labels and columns the predicted ones reports the inference performance achieved by the classifier (Fig.5d). We found that cross-validated classification performance was poor (mean accuracy for all conditions upon cross validation trials ∼18%; Fig.5d, 5g; see also Fig.S8). Therefore, we could not extract sufficient information from synaptic weights distribution in microzones to sort connectivity maps between conditions.

We then described the fine structuration of connectivity maps in terms of graph-based metrics. Unlike microzone synaptic weights, graph metrics have a complex dependency on map details at many different scales simultaneously and can thus transcend anatomical subdivisions. In this alternative description, we included similar number of graph-based features as in the aforementioned descriptions in terms of microzone synaptic weights: the whole-map modularity and lateralized (ipsi vs contralateral nodes) participation, degree z-score and assortativity. We then compared these features across conditions. For all groups and for all features we found a large variability across different maps and specimens (Fig.S7). Such large within-group variability of each individual feature prevented us from detecting any significant group-level difference between the different adaptive conditions and the CTRL maps (Fig.S7). Nevertheless, these individual graph metrics carried useful information for condition discrimination when considered together. A t-SNE projection based on these alternative vectors of graph-based metrics showed that the cloud of maps for the different conditions were displaced with respect to the CTRL maps (Fig 5c) and that LS/ES and LC/EC clouds were shifted away from the cloud of the STR/LTR conditions. Furthermore, our supervised machine learning classifier now yielded a cross-validated classification performance larger than with synaptic parameters and well above chance level (Fig.5c, e). In particular, ∼80% of EC and ∼70% of ENR maps were correctly classified. The largest correspondence occurred between maps in EC condition, whereas the mean accuracy for all the subtypes across cross validation trials was >60% (Fig.5e). See Fig.S8, for coarser (global) or finer (microzone-level) graph parameterizations which do not further improve the performance. Taken together, these findings suggest that while connectivity maps varied between animals in each condition, the reproducible correlation between some microzones (Fig. 5b) and the fact that the locomotor condition in which a map was obtained can be inferred to a certain extent from the graph descriptors (Fig. 5d, e) suggest that the network structure exhibits invariant features for each of these behavioral conditions.

### Individual connectivity maps reflect individual-specific behavioral features: map individualities

The large variability between maps may be due to experimental “noise” or reflect on the contrary fine levels of behavioral differences across individuals, rather than adaptive conditions. To investigate this hypothesis, we tried predicting individual-level specificities in adaptive locomotor behavior from graph descriptions of the global maps and behavioral performance either on the wheel (TR) or on the force pressure corridor (LS, LC). We used the multi-dimensional graph-based map vectors and trained generalized linear models (GLMs) to predict individual-level behavior from spatial organization of their maps. In the TR condition, an individual mouse had a short (7 days) or long (19 days) training in the running wheel yielding different total cumulative distance performed more or less eagerly (as quantified by the slope of distance travelled vs days trajectory, Fig 3a). Figures 6a and 6b show the correlation between, respectively, the actual values of total training distance/training slope (Fig3a, and the corresponding values predicted by a generalized linear model trained on graph-based map vectors (methods). The cross-validated generalization performance was significantly above chance level for both features (insets of Figures 6a and 6b), reaching an actual-to-predicted R-values fits of 0.64±0.06 (actual) vs -0.02±0.07 (shuffled, CI=95% permutation testing) for total distance and 0.37±0.06 (actual) vs 0.04±0.07 (shuffled, 95% CI, permutation testing) for motor training slope.

**Figure 3.**
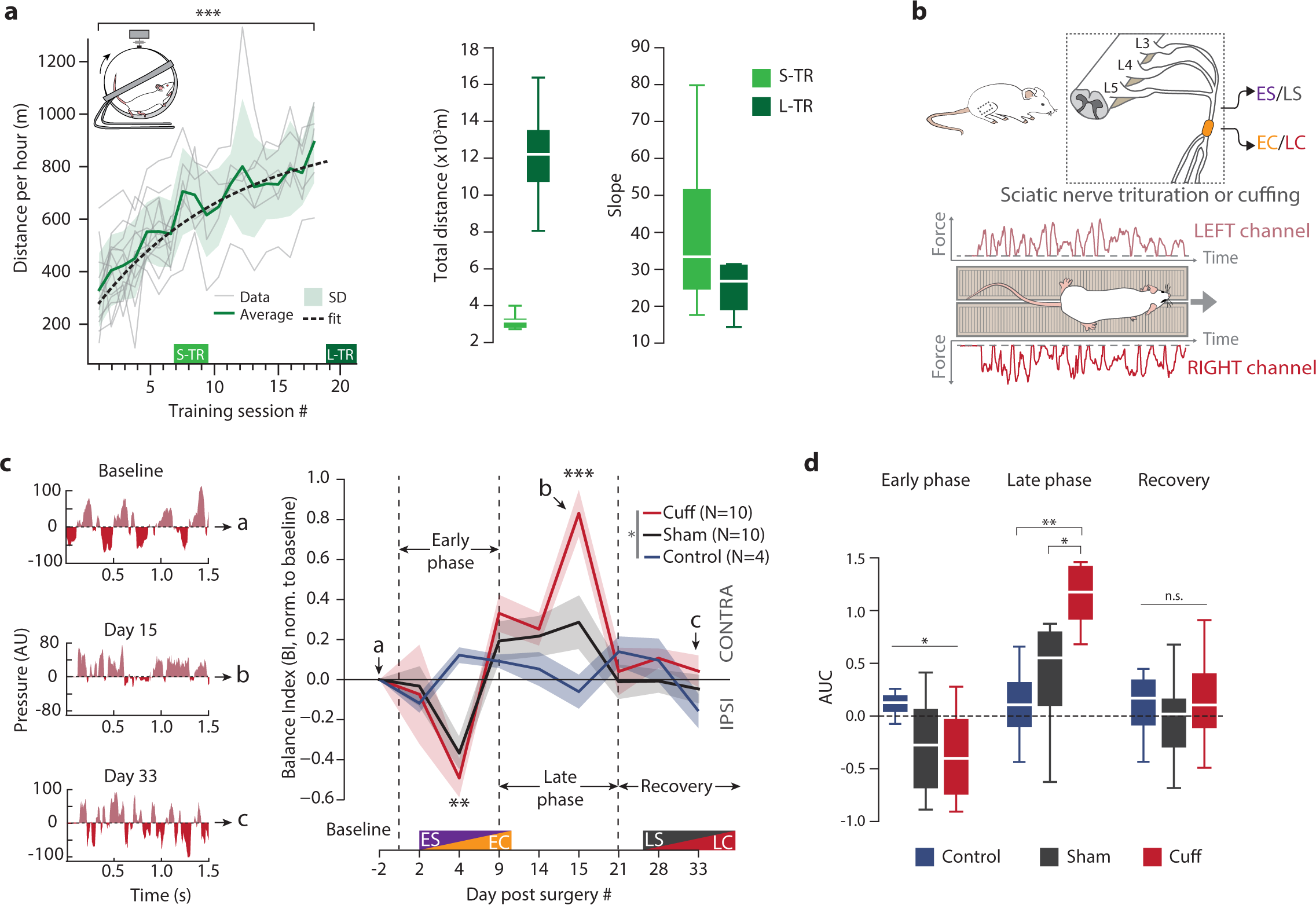
Locomotor adaptation in different contexts. **(a)** *Left panel*, Overall performance of mice in voluntary locomotor training in a wheel (1h/day for 7 days in S-TR and 19 days for L-TR, average±SD). ***: paired t-test between session1 and session19, p=2.10-4. Dashed line, logistic fit, p=2.75.10-26. ***Right panels***, total distance, and slope of the distance across sessions for S-TR and L-TR groups. **(b)** *Upper panel*, cuff model of locomotor impairment. In EC and LC groups, a polyethylene cuff is surgically wrapped around the main branch of the right sciatic nerve. In sham (ES, LS) no cuff was inserted but the sciatic nerve was triturated. ***Lower panel***, balance was evaluated on a corridor mounted with force sensors and pressure from left and right limbs was measured during each trial. **(c)** Balance deficits in cuffed and sham groups. ***Left panel***, examples of raw data measurements from force-sensor corridor in a cuffed mouse before surgery (*baseline, a*), 15 days after surgery (*b*) and 33 days after surgery (*c*). Left/right body side is shown in light/dark red. Balance Index (BI) is computed as the ratio of the integral of the signal from the left and right side. ***Right panel***, averaged (±SEM) time-course of the balance index in cuffed (LC), sham (LS) and CTRL animals normalized to baseline. *Early phase*, from day 0 to day 9 post surgery; *late phase*, from day 9 to day 21 post surgery; *recovery*, from 21 post surgery. One-way repeated measurements MANOVA from day 2 to day 28, *p=0.0433; post hoc ANOVA, **Cuff vs Ctrl vs Sham ANOVA, p=0.0044; ***Cuff vs Ctrl vs Sham ANOVA, p=0.0025. (d) Area under the curves (AUC) for cuffed, sham and CTRL groups. Levene’s test, *pEarly= 0.0205, pLate=0.7264, pRecove- ry=0.91852; Kruskall-Wallis followed by post hoc Mann Whitney U tests, pKruskal-Wallis-Early=0.2122; pKruskal-Wallis-Late=0.02107; pKruskal-Wallis_Re- covery=0.644; pMWU_CTRLvsSHAM_Late=0.1444; *pMWU_SHAMvsCUFF_Late=0.02257, **pMWU_CUFFvsCTRL_Late=0.009812.

**Figure 4.**
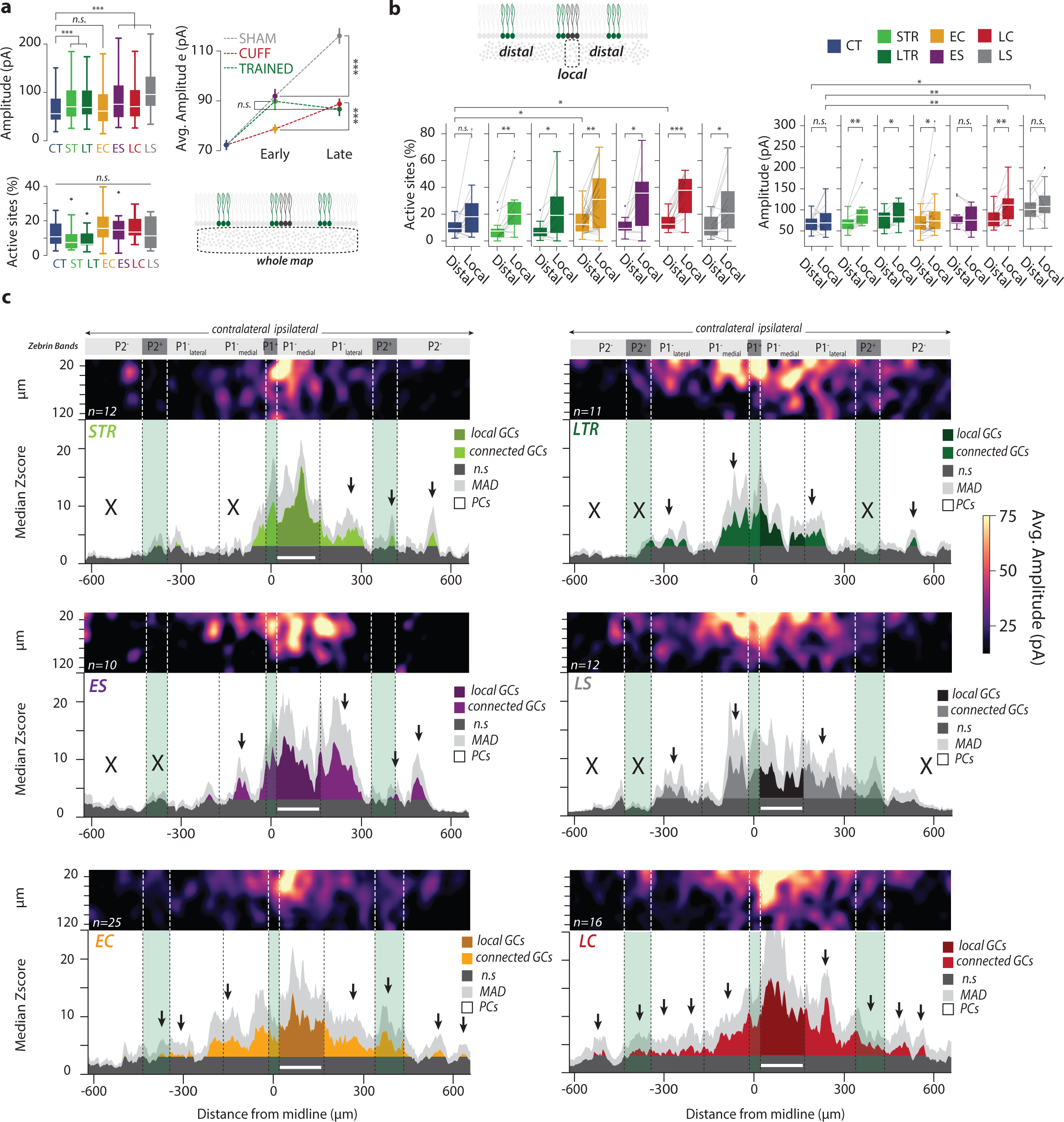
Synaptic adjustments of GC-PC maps following motor adaptation. **(a) *Top left***, distributions of synaptic amplitudes of significant sites recorded in each behavioral condition. For statistics, see table in Figure S10. ***Top right***, average synaptic weights at different time points of locomotor adaptation. Sham (in grey, ES and LS pooled); CUFF (in red, EC and LC pooled) and TR (in green, STR and LTR pooled). For statistics, see table in Figure S10. ***Bottom***, proportion of active sites in maps. Kruskal-Wallis test, p=0.12681. **(b)** Percentage of active sites and average synaptic amplitudes measured in local (*0-120 µm from midline*) vs distal GCL. Results of statistical comparisons are available in Figure S10. **(c)** 2D averaged synaptic connectivity maps (***top***) and 1D median patterns (***bottom***) in each locomotor condition. The white bar illustrates PC positions. GC sites in dark grey are silent. Black crosses/arrows highlight specific silent/connected areas. Light gray curve represents Median Absolute Deviation (MAD).

**Figure 5.**
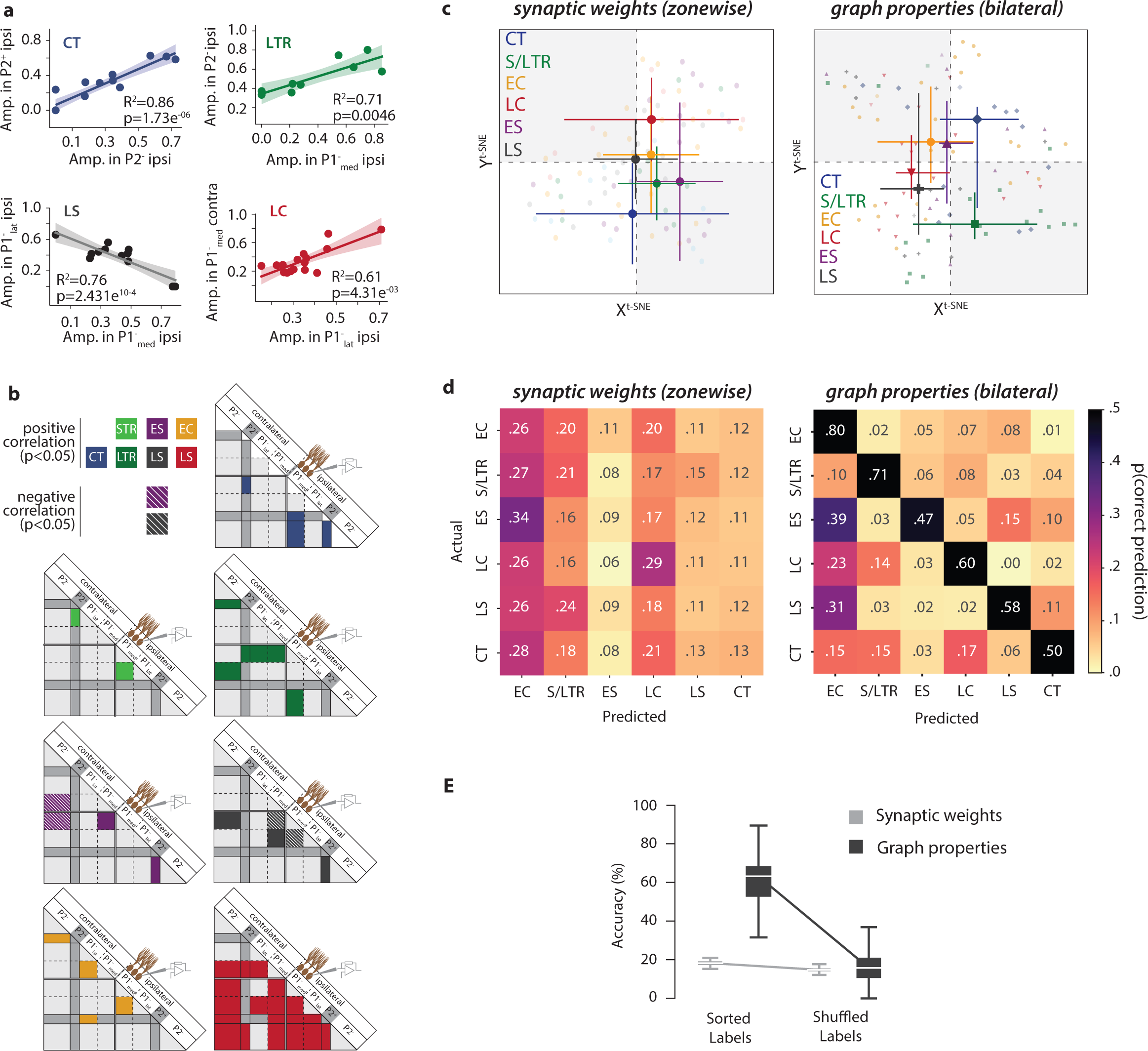
Graph features of synaptic maps can predict locomotor conditions. **(a)** Examples of microzone-wise Pearson correlations of normalized averaged synaptic weights in connectivity maps. Shaded curves represent confidence interval. **(b)** Diagram illustrating Pearson correlation between microzones in each locomotor condition. Colored squares indicate significant correlations (*p<0.05*) while dashed squares represent significant anti-correlations (*p<0.05*). **(c) *Left panel***, t-SNE representation of connectivity maps based on averaged synaptic weights by microzones. Individual synaptic maps were defined as a vector composed of the average synaptic weight in each microzone. This 8-dimensional space has been reduced via t-SNE to be visualized in a 2D space. Individual dots represent the position of each map in the t-SNE space. Big dots represent supervised cluster median, and error bars show .75 median quartile. Right panel, t-SNE representation of maps based on graph properties. ***Right panel***, Identical analysis as in *(c, left panel)* but with graph-based vectors. **(d) *Left panel***, classification of synaptic maps based on synaptic weights. Map vectors described in *(c)* were used to train a supervised random forest classifier. Accuracy of classification is displayed as a confusion matrix, where values show the probability (from 0 to 1) to sort a map with actual label (*right*) to predicted category (*down*). Right panel, classification of synaptic maps based on graph properties. ***Right panel***, Identical analysis as in *(d, left panel)* but with graph-based vectors. **(e)** Average accuracy of random forest classifier. Accuracy of the classification with synaptic parameters or graph properties in each cross-validation iteration were pooled and shown here as box plots. Random forest classification was done either with exact label correspondence (i.e., map vector is correctly labeled, *“Sorted labels”*) or with shuffied labels (i.e., each map vector has been given a random label among CT/TR/ES/EC/LC/LS groups).

**Figure 6.**
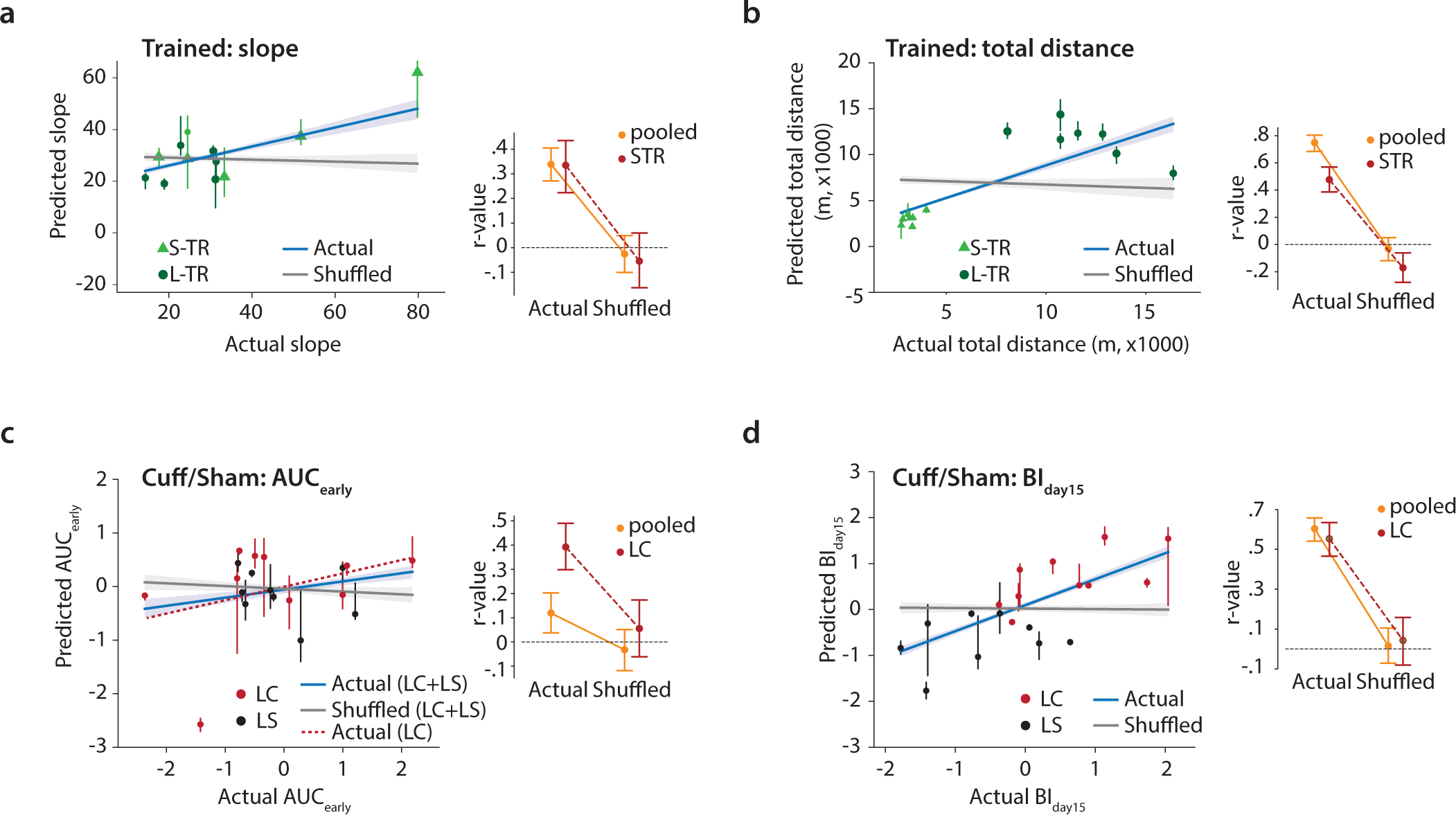
Graph properties of connectivity maps predict behavioral individualities. Results of behavioral features prediction based on graph properties of synaptic maps using Generalized Linear Models (GLMs). In TR animals, the slope of performance **(a)** and the total travelled distance **(b)** on the running wheel were considered for prediction. In LC and LS groups, post-surgery imbalance (*AUC_early_*, **c**) and maximal impairment (*BI_day15_*, **d**) were considered for prediction. In every plot, average predictions are scattered on left panels, and regression coefficients (r values) are plotted in right panels. Linear models with actual data are shown in blue while results with shuffled data (i.e., chance level) are shown in gray.

We then considered the LC and LS groups, for which it was possible to follow the locomotor recovery following surgery across several weeks. We used the AUC_early_, AUC_late_ and BI_day15_ features (see Fig. 3c, d) as possible prediction targets. The prediction of the early component AUC_early_ (Fig.6c) was only marginally significant (cross-validated R-value fit of 0.15±0.08 vs - 0.05±0.08 for shuffled; 95% CI, permutation testing), even if it raised to 0.33±0.1 vs -0.06±0.12 for shuffled (95% CI, permutation testing) when limiting the analysis uniquely to LC specimens. The prediction of the late component (AUC_late_) gave similar poor results (Fig. S9). Nonetheless, we were able to significantly predict the evolution of locomotor unbalance monitored by the BI_day15_ component from graph description characterizations of connectivity maps (Fig.6d), with a cross-validated R-value fit of 0.65±0.05 vs 0.01±0.1 for shuffled (95% CI, permutation testing).

We thus conclude that the characteristic variability of the fine spatial structure of GC-to-PC connectivity maps is not mere “noise” but rather reflects specific overtraining or post-traumatic recovery adaptation at the single individual level.

## Discussion

### Multi-microzonal connectivity maps adaptation to locomotor context in the cerebellar cortex

We identified patchy GC connectivity maps to medial PCs as observed in our previous study^14^. These maps are deeply modified when the locomotor context is modified and during the PND14-PND18 developmental stage. Spatial synaptic analyses were based on CF/MF inputs topography ^32, 42, 47, 57^ (Fig.S1). In CTRL animals, connected patches were specifically found in zones receiving information from the distal and proximal hindlimbs either from lumbar and thoracic spinocerebellar (P2^+^/P1^-^_lateral_ and P1^-^/P1^+^_medial_) or from ponto-cerebellar MFs (P1^+^ and P2^+^)^14, 47, 48, 57–61^. Conversely, central P1- patches which are mainly innervated by the external cuneate nucleus and mediate forelimb proprioceptive inputs are rarely connected. These results indicate that medial PCs integrate both somatosensory and cerebral MF information from both hindlimbs. In TR and perturbed conditions (sham and cuffed), we observed an increase in synaptic weights and context specific spatial synaptic reorganization of connected GC patches, particularly in the ipsilateral side of the GCL. While in CTRL condition connectivity maps were symmetrical (Fig.1e), in cuffed (EC/LC) and TR conditions symmetry vanished and synaptic weight as well as percentage of active site increased in the ipsilateral part (Fig.4, S3). Moreover, in invasive conditions (cuffed/sham) the asymmetry increased after full recovery (EC/ES vs LC/LS). These results not only suggest that synaptic plasticity remodel connectivity maps as shown already in^14^ but also locomotor conditions requesting intense re-adaptation such as LC recovered with denser connectivity maps including new GCL areas receiving MFs from both forelimbs and hindlimbs (e.g. P2^-^, Fig S1 and Fig4). We postulate that P2^-^ microzones which target hindlimb extensor muscles^29, 62, 63^ are more involved in postural control in cuffed and TR animals as an intense reorganization of the cooperation between forelimbs and hindlimbs is required^64^. Hence, medial PCs in LC animals have extended receptive fields to both limbs as shown *in vivo* after electric parallel fiber stimulation in the molecular layer^65^. Such structuration is in agreement with the fractured somatotopy identified with micromappings^66^ and recent imaging studies showing dense GC activation upon activity^67–69^. We suggest that the fractured somatotopy of the MF inputs underlie the topography of the context specific reorganization of connectivity maps.

*In vivo* and in vitro experiments suggested that PCs have a high probability of connection with local GCs^70–72^. We confirmed these results (Fig.4) as local GC sites were strongly connected in all individual maps and all contexts suggesting that they provide a non-conditional input. During development, we observed that strong connections arose from local GCs although PFs are already hundreds µm long (Fig.S5a). It has been shown that local GCs are unable to elicit LTD through their ascending axon^73^. We suggest that MF inputs always significantly influence PCs belonging to their incoming microzone, indicating that the topography of MF inputs predict a minimal arrangement of coordinated microzones during movement as it has been recently observed for CF activated microzones^31^. Therefore, MFs conveying information from hindlimbs and forelimbs which project to multiple microzones (Fig.S1) are likely to activate medial PCs as well as PCs in the other microzones in a coordinated manner. We further demonstrate here that GCs will favor and refine this population code in a context dependent manner yielding a timely organized sequence of activations. This spatial arrangement of multi-microzonal communication may define a functional module dedicated to a specific adaptive behavior in a given locomotor context^41^.

### Graph networks identified a critical period

Graph-based modules which describe the functional structuration of GC areas were built by aggregating correlated columns of GCL in an unsupervised manner (Fig.1 and methods), without *a priori* on the map spatial organization. As such, the features of the graph community structure can account simultaneously for the degree of patchiness and for the tight links between distant GC patches, being thus able to account for the complexity of MF somatotopy. Graph networks are a powerful tool in the description of functional microcircuit organizatio (*see below*). For example, during postnatal development, we observed that between P14 and P18, connectivity maps were highly variable with a median 1D pattern similar to randomized organization^74^. Synaptic weights were larger, and the proportion of active site increased by 2- fold in the upper part of the GC layer. Graph parameters allowed us to quantify and highlight significant changes in the graph encoding of synaptic maps, as modularity and participation dropped to minimal values indicating that modular structuration vanished at this stage before resuming to higher values in adulthood. Between P14-P18, mice start to walk and open their eyes^52, 53, 75^. They are hyperactive with bouts of vigorous jumping called *‘hoppy’* or *‘popcorn’* stage resulting from synchronous contraction of fore and hind limb extensors^53, 76^. Moreover, several important molecular and morphological modifications occur at this stage. For example, the GluN2C subunit incorporation in GC NMDA receptors leading to an enhancement of GC excitability^77^ and the regression of CF multi-innervation allowing proper LTD induction at the GC-PC synapses^78^. Therefore, we postulate that the P14-P18 period is a critical period allowing thorough reorganization of connectivity maps as observed in the visual cortex^79^.

### Capturing the shape of map organization

Since connectivity maps are not only non-randomly organized, but that their organization is highly and specifically adaptive, it is therefore a question of utmost importance to understand how to appraise the spatial complexity of this maps’ adaptive organization into a few quantitative metrics. Here, we designed a mapping from 2D maps to graph descriptions and described topological features of synaptic maps. The pertinence of our mapping can be validated post hoc by the superiority in prediction that descriptions in terms of graph metrics confer with respect to other descriptions apparently more tightly linked to physiology (Figures 5d, e). Such superior performance may be linked to the fact that the fine parameterizations of connectivity correlations used, capture aspects of network organization which are neither purely local (depending on a single map location), nor exclusively global (properties common to all map locations) as for example the redundancy of specific MF inputs in lobule IV (Fig.S1). In particular, even when they are site-specific and heterogenous across map locations, graph features describe properties of the coupling of these locations with many other locations. They thus reflect structure at many different and nested intermediate scales^80^, that overlap only imperfectly with traditional anatomical subdivisions and that reveal, in a sense, a mixture of “segregation” and “integration”, two notions more commonly invoked in reference to neuroimaging data^81^. As a matter of fact, individual-specific adaptive behavior can be predicted by graph features even when averaged over all the sites in the map, thus fully ignoring anatomy (Fig.6). At the same time, the discrimination between conditions is improved by using lateralized (ipsi vs contra) with respect to global graph metric averages but averaging to the level of individual microzones does not yield additional improvement (Fig.5d vs Fig.S8). It is thus likely that the number of available data for classifier training puts a limit to the amount of information that can be extracted without overtraining^82^ and that more precise characterizations could be still achieved by using larger datasets.

We remark that classification and prediction of adaptation behavior is necessarily *polythetic*^83^, i.e. require to simultaneously monitor correlations between a multiplicity of graph features, since features taken one at a time have a dramatic variability and largely overlapping ranges between classes, as happens also e.g. for neuronal types^84^. Different adaptive conditions thus give rise to broad phenotypes of maps, in which connectivity is, nevertheless, not deterministically constrained, but preserves a large freedom, exploitable to give rise to accurate behavior adjustments. Graph-based features are only a first step toward the quantitative characterizations of the connectivity maps specificities at the single individual level. In the future, more powerful and general topological data analysis approaches^85^ may be used to capture the own “shape” of each map and how the individualities of this shape reflect individual behavioral histories^86^.

### Individualities: implications for internal model storage

Many computational models suggest that the cerebellar microcircuits learn internal models of the body that are designed to predict an expected sensory feedback of a given command or simply update a motor command as suggested by many models^17, 20, 21^. A change in the relationships between muscles or a modification of limb alternation during locomotion as we have performed in our experiments should affect the workflow of computation involving internal models in the cerebello-thalamo-cortical loop as also suggested in^87, 88^. We here provide strong evidence that PCs learn to adapt locomotor behavior by adjusting GC-PC connectivity maps in a context dependent manner. In medial PCs, each context (e.g., gallop vs walk or sham vs cuff) lead to connections with a specific set of microzones through GCs. Moreover, multidimensional analysis of graph parameters (Fig 5 and 6) defines a mathematical fingerprint of each connectivity map which is predictive of individual behavioral features. We therefore argue that the specific distribution of GC sites associated with medial PCs in the different behavioral conditions relates directly to locomotor adaptation and internal model adjustments.

## Acknowledgements

This work was supported by the Centre National pour la Recherche Scientifique (CNRS), the Université de Strasbourg, the Agence Nationale pour la Recherche (ANR-2015-CeMod, ANR-2019-MultiMod, ANR-2019-NetOnTime) and by the Fondation pour la Recherche Medicale to PI (# DEQ20140329514). JB and DB acknowledge support by the Agence Nationale pour la Recherche (SENCE, ANR-18-CE37-0018-03). LS was funded by fellowships from the Ministère de la Recherche and Fondation pour la Recherche Médicale. We thank Nathalie Petit-Desmoulières, Dr Elise Savier, Dr. Sophie Reibel-Foisset and the staff of the animal facility (Chronobiotron, UMS 3415 CNRS and Strasbourg University) for technical assistance. We thank Antoine Valera, Matilde Cordero-Erausquin, Didier Desaintjan and Frédéric Doussau for critical readings.

## Methods

### Ethics & Animals

All experiments were conducted in accordance with the guidelines of the *Ministère de l’Education Supérieure et de la Recherche* and the local ethical committee, the *Comité Régional En Matière d’Expérimentation Animale de Strasbourg* (CREMEAS) under the agreement n° A67-2018-38 (delivered on the 10th of December 2013 to the animal facility Chronobiotron UMS3415). 83 AldoC-venus^45^ male mice from PND9 to PND60 under CD1 background were used for this study. Mice were housed by 3 or 4 littermates per cage, in conditions required to fulfil their ethogram with food and water *ad libitum* in a 12/12 light/dark cycle.

### Surgical procedures for Cuff and Sham mice

Mice were anesthetized by inhalation of isoflurane (Verflurane, Virbac, France, 4% for induction then 1-2% for the surgery). Mice were laid at rest on the left side of the body to expose the right hindlimb. A mix of lidocaïn/bupivacaïne (2 mg/kg each) was subcutaneously injected prior to incision. A 0.5 cm incision was made parallel to the femur to expose leg muscles. Muscles were gently separated using sterilized wooden sticks to expose the main branch of the sciatic nerve. The nerve was pulled out of the limb and a sterile 2 mm section of split PE-20 polyethylene tubing (cuff), 0.38 mm ID / 1.09 mm OD, was wrapped around the nerve with the help of a pointed steel stick and a bulldog clamp. The nerve was then pushed back under the muscle fascia, and skin was sutured^91^. For sham mice, the same procedure was followed except that no cuff was implanted. After surgeries mice received an intraperitoneal injection of non steroidal anti-inflammatory drug ( Metacam, 2 mg/kg) and were left at rest for 24h minimum before behavioral assessment. In cuffed animals, the plastic cuff remained around the sciatic nerve until the sacrifice of the animals.

### Monitoring Balance

To assess and quantify cuff-induced gait and balance impairments during locomotion, we built a force-sensor device. Mice were trained to walk in an 80 cm-long corridor covered with two parallel strip-shaped force-sensors on each side of the corridor. The width of the corridor and the space between the two strips were adjusted to ensure that left and right limbs of CD1 mice activate the same strip. Force sensors (FSR, 10x622mm each, FSR 408, Interlink Electronics, USA) are two-wire devices with a resistance that depends on applied force, allowing simple force-to-voltage conversion when tied to a measuring resistor in a voltage divider. As mice enter the corridor, force and video acquisition were triggered by a Raspberry Pi microcomputer (Raspberry Pi Foundation, UK) connected to an IR-barrier. Voltage-divider output was linked to an Analog-Digital interface (NI USB 62-11, National Instruments, USA). Force signals for the left and right side of the body were acquired simultaneously using WinWCP 4.2.2 freeware (John Dempster, SIPBS, University of Strathclyde, UK). Recordings were digitized at 15-20 kHz.

The balance index (BI, Fig.3) corresponds to the ratio of the integrated force-signal from each side of the body. All recordings contained at least 5 consecutives strides, otherwise they were discarded. or further technical details (i.e., apparatus dimensions, scripts and wiring diagrams) see https://github.com/ludo67100/cerebellarMaps/tree/main/Balance_supplementary

In cuffed (EC/LC), sham (ES/LS) and CTRL animals, the balance index was determined before and after surgeries every 2-3 days for 1 month. A time-course was established, and we estimated the cumulative gait imbalance during the early days (AUC-early for area under the curve; see Fig.3), late phase (4-21 days after surgery; AUC-late) and maximum imbalance (balance index on the 15th day after surgery).

### Locomotor training in a wheel

TR group of Mice had access to a running wheel for 1h/day during 7 (S-TR group) or 19 (L-TR group) consecutives days. For each session, mice were removed from their housing cage and placed in another individual cage equipped with a vertical, access-free running wheel. Locomotor activity was monitored using a piezoelectric sensor on the cage coupled to a magnet on the wheel to count the number of wheel turns during each session. We measured the total distance covered during the training period of 7 or 19 days and the slope of the training trajectory using linear regression analysis.

### Di-I injections to measure parallel fiber extension

PND8-9 CD1 pups were placed in crushed ice for 2-3 minutes to be anesthetized. A small incision was rapidly made over the cerebellum and .5 to 1µL fluorescent dye (Vybrant DiI cell labeling solution, Thermo Fisher) was injected at the midline of lobules IV/V of the cerebellar cortex using a glass pipette and a pressure pump (Picospritzer III, Parker, USA). Location and depth of the injection were determined by eye using visual cues. After injection, the opened skin was closed with a drop a biocompatible glue (Vetbond, 3M, USA) and pups were placed on a heat pad a few minutes prior return to the mother in the homecage.

PND30 CD1 mice were anesthetized by inhalation of isoflurane (Verflurane, Virbac, France, 4% for induction then 1-2% for the surgery) and mounted on a stereotaxic frame (Model 68526, RWD Life Science). Body temperature was monitored using a rectal probe and maintained with a heat pad. A mix of lidocaïn/bupivacaïne (2 mg/kg each) was subcutaneously injected over the skull prior to incision, followed by an intraperitoneal injection of a non-steroidal anti-inflammatory drug (Metacam, 2 mg/kg). A parasagittal incision was made over the skull to expose lambda and bregma landmarks. The skull was cleaned using cotton sticks soaked in sterile 0.9% NaCl solution (saline). A 0.5 mm diameter hole was drilled at AP =-2mm, ML = 0 (from Lambda) to expose lobules IV/V. DiI was injected as described for pups. Skin was sutured after injection and animals were put back in their home cage

### Slice preparation for electrophysiology and photostimulation

Slices were prepared from P9–P90 male CD1 ALDOC mice. P12 to P90 Mice were anesthetized by inhalation of isoflurane 4% (Verflurane, Virbac, France) and then killed by decapitation. P9 and P10 pups were sedated by hypothermia prior to decapitation. The cerebellum was rapidly dissected out and placed in ice-cold (<4°C) artificial cerebrospinal fluid (ACSF) continuously bubbled with carbogen (95% O_2_, 5% CO_2_), containing (in mM): NaCl 120, KCl 3, NaHCO_3_ 26, NaH_2_PO_4_ 1.25, CaCl_2_ 2.5, MGCl_2_ 2, glucose 10 and minocycline 0.00005 (Sigma- Aldrich, USA). 300 µm-thick transverse acute cerebellar slices were then prepared (Microm HM 650V, Microm, Germany) Fig.in ice-cold (>4°C) N-methyl-D-aspartate (NMDG) based solution containing (in mM) : NMDG (93), KCl (2.5), NaH_2_PO_4_ (1,2), NaHCO_3_ (30), HEPES (20), Glucose (25), sodium ascorbate (5), Thiourea (2), sodium pyruvate (3), N-acetylcysteine (1), Kynurenic acid (1), MgSO_4_7H_2_O (10), CaCL2.2H2O (0.5). After cutting, slices were maintained in 34°C ACSF at least 45 min, then kept at room temperature until the end of the experiment.

### Patch clamp recordings

Whole-cell patch-clamp recordings in voltage-clamp mode were obtained using a Multiclamp 700B amplifier (Molecular Devices, USA) and acquired with WinWCP 4.2.2 freeware (John Dempster, SIPBS, University of Strathclyde, UK). Patch pipettes (3-4MΩ) were pulled with borosilicate capillaries (Warner instruments) using a gravitational puller (model PC12, Narishige, Japan). Series resistance was monitored and compensated (70%–80% typically) in all experiments, and cells were held at -60 mV to isolate excitatory postsynaptic currents (EPSCs. The internal pipette solution contained (in mM): CsMeSO4 135, NaCl 6, HEPES 10, MgATP 4 and Na_2_GTP 0.4. pH was adjusted to 7.3 with KOH and osmolarity was set at 300 mOsm. Biocytin (Sigma Aldrich) or neurobiotin (Vector Laboratories, USA) were added (1 mg/ml each) for cell reconstruction. Voltages were not corrected for the liquid junction potential, which was calculated to be 9 mV (i.e., the membrane potential was 9 mV more hyperpolarized than reported). We accepted recordings for which the inward current at -60 mV did not exceed 1 nA. Synaptic currents in PCs were low pass filtered at 2.4-2.6 kHz, then sampled at 20 to 50 kHz. All recorded cells were in lobule III or IV. All experiments were performed at room temperature using the same bubbled ACSF than for dissection. We systematically blocked NMDA, adenosine, CB1, GABAB and mGluR1 receptors to limit the modulation of EPSCsamplitude by activity-dependent activation of these receptors. They were respectively blocked using (in mM): D-AP5 0.05 (Ascent Scientific, Abcam Inc), DPCPX 0.0005, AM251 0.001, CGP 52432 0.001 and JNJ16259685 0.002 (Tocris-Cookson, UK). For excitatory maps only, inhibitory transmission was blocked with Picrotoxin (0.1mM).

### Photostimulation

Uncaging experiments were performed using RuBi-Glutamate (100µM) perfused in the recording chamber in a closed circuit (Abcam, UK). To map GC-PC synaptic connections, slices were positioned according to the horizontal plane in the recording chamber. In ALDOC mice^45^, Venus fluorescence allowed precise visualization of PCs of the P1^-^_medial_ zebrin band. A micromirror DMD device control by the IQ3 software (Mosaïc, Andor Technology, Belfast, Ireland) mounted on an optic microscope (Olympus BX51, Japan) allowed systematic blue light photostimulation (pulses of 30 ms) of 20*20µm or 40*40µm squared GCL regions with LED- (460nm, Prizmatix, Israel) through a 40X objective (Zeiss, Germa (Fig.1B). The developmental dataset was performed with a grid of 128 sites (40µm² per site; low-resolution) while synaptic maps recorded after locomotor adaptation were obtained with a high-resolution photostimulation grid containing 384 sites (20µm² per site). A single grid covers 320 µm of the GC layer in the mediolateral axis. However, one synaptic was composed of 4 fields of view covering up to 1230 µm in the mediolateral axis. Two stimulations of the same GC site were separated by 60 seconds minimum. For each experiment, a single site of the granular layer was photostimulated between 5 and 10 times in total (yielding 5 to 10 recordings for averaging and analysis).

### Immunohistochemistry

After recordings, the patch pipette was gently removed from the recorded PC and the slice was immediately transferred from the recording chamber to a fixation solution composed of 4% paraformaldehyde (Electron Microscopy Sciences) in ACSF for a maximum of 24h (4°C).

Zebrin bands were identified using intrinsic venus fluorescence. Recorded cells were labeled using Alexa 555-Streptavidin (Thermo-Fisher, 1/1000, 3 hours at room temperature). Ipsi- and contralateral P1+, P1-, P2+ and P2- Zebrin bands and distance between the recorded PC and the midline (P1+) were measured in each experiment. In adult CD1 mice, averaged Zebrin band lengths in lobule III/IV were (in µm±SD) : *P2- contralateral* 416.6±70.72; *P2+ contralateral* 71.56±24.59; *P1- contralateral* 320.516±62.94; *P1+* 34.63±16.18; *P1- ipsilateral* 320.46±60.54; *P2+ ipsilateral* 69.75±22.53; *P2- ipsilateral* 438.04±64.25. Recorded PCs were located at 52.42±29.8 µm from the midline of lobules III/IV, corresponding to the cluster 1 defined by Valera et al.^14^.

### Data processing and analysis of synaptic connectivity

Data processing and analysis of synaptic parameters were performed via homemade scripts and routines written in Python 3.6 using the following packages : Pandas, Scipy, Sci-kit learn, Numpy, Matplotlib, Neo 0.8, Orange Data Mining^92^. Multivariate analysis (one way repeated Manova) was performed using the Real Statistics Resource Pack software (Release 6.8). www.real-statistics.com. Python-based code is available at: https://github.com/ludo67100/cerebellarMaps

### Reconstruction of 2D connectivity maps and 1D input patterns

In each GC site of the grids, EPSCs elicited by RuBi-glutamate uncaging were measured in a 200ms time window from stimulation onset (A_stim_, Fig.S2B). Each GC sites was photostimulated 5-10 times and EPSCs were averaged. For comparison, spontaneous activity in the slice was measured on each averaged recording of the map as the minimum amplitude in a 200ms window 1s before or after the photostimulation (A_noise_, Fig.S2B). Histogram of spontaneous activity was then fitted with a gaussian function and the standard deviation (σ) of the synaptic noise was used to estimate z-scores of EPSCs (Fig.S2B).

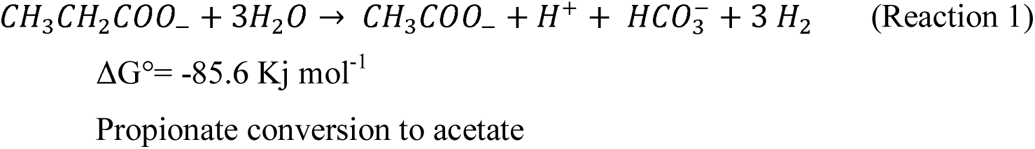

A zscore > 3.09 corresponds to a significant synaptic with an alpha=0.01. Z-scores were then used to draw 2D synaptic maps (Fig.1). For each GCL column, we used the maximum z score value to define a 1D projection pattern (Fig.1).

### Median input pattern and averaged synaptic maps

In each behavioral condition, we built the median 1D GC-PC inputs patterns and the averaged 2D synaptic maps. Individual 1D input patterns (described above) were interpolated and convolved with a triangular kernel (width=9µm) prior to median computation. For averaged 2D maps, individual maps and corresponding positional arrays were concatenated in separate vectors. The positional vector was sorted in ascending order along the mediolateral axis (from contralateral to ipsilateral positions in the GCL) and the exact same sorting rule was applied in parallel to the map-based vector. The resulting, spatially sorted meta-map vector was then divided in 30µm-wide bins for averaging, yielding average maps shown in Fig.1, 2 and 4.

### Synaptic weights correlations between microzones

In each map, synaptic weights from GC sites were normalized to the maximal amplitude. Amplitudes from significant GC sites (i.e., Z score > 3.09) were averaged by microzonesand Pearson correlation between microzones were computed in each condition (Fig.5A).

### Graph properties

#### GC-PC maps pre-processing

We considered each two-dimensional connectivity map as a matrix ***M***, whose *M_xy_* entry denotes the EPSC amplitude recorded at the position coordinates (x,y). We then focused on the column vectors ***M***_*x*_ of this matrix and for every pair of positions *x* and *x’* along the scanned mediolateral axis we computed the normalized Pearson correlations *C_xx_’ =* corr(***M***_*x*_, ***M***_*x’*_) between the connectivity profiles at the two considered positions. *C_xx_’* values were then compiled in a correlation matrix ***C****(**M**)* (*“Raw matrix”* on Fig.1D) that captured the patchy structure of the original map. ***C****(**M**)* can be considered as the adjacency matrix of an undirected graph. The entries *C_xx_’* thus become the strengths of links between graph nodes associated to the positions *x* and *x’*. The more similar are the connectivity profiles between two positions *x* and *x’* and the stronger will be the weight *C_xx_’* of the connection between them. A standard Louvain algorithm^51^ is then used to optimally partition graph nodes into non-overlapping communities such that total weight of within group edges are maximized and weight of between group edges are minimized. A resorting of node labels according to the extracted community labels leads to the “Rearranged Graph Matrix” in Fig.1D in which a block structure is better visible than before reordering of nodes.

#### Graph-based features

All the graph properties were calculated using the *Brain Connectivity Toolbox* for python (https://pypi.org/project/bctpy/) designed by M Rubinov and Olaf Sporns^50^

We characterized the GC-PC maps using graph properties such as:

a. Modularity index - Modularity index maximizes the number of within group edges and minimizes the number of between group edges using community the Louvain algorithm. It is a measure of how modular or “patchy” a graph is, i.e. higher the modularity index, higher is the number of modules or “patches” in a graph. Modularity index is a graph centric measure and hence we get one value per map/graph. We use a variant of modularity index measure designed for weighted graphs given by the following equation (Newman 2004, Rubinov & Sporns 2010):

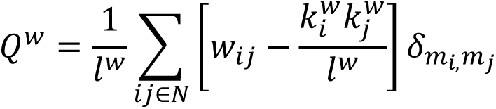 Where *l*_*w*_= total weight of the graph; *W*_*ij*_ = weight of the edge between node *i* and node *j*; 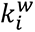, 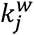 = weighted degrees of node *i* and *j* respectively; 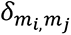 = 1 if *m_i_* = *m_j_* and 0 otherwise; *m*_*i*_,*m_j_* are the modules containing nodes *i* and *j* respectively.
b. Module degree z-score (weighted) – is a measure of within-module version of degree centrality. Degree centrality is a measure of edges connected to a node. Hence, a weighted module degree z-score measures how strongly is the node connected to other nodes within the module. A high average module degree z-score for a correlation map indicates that patches in the map tend to be organized around a center, fading in a graded manner into the background which does not elicit stimulation responses in the considered PC. On the contrary, maps with lower average module degree z-score have internally homogeneous patches with sharper edges. Module degree z-score is a node centric measure i.e, we get a value per node. The module degree z-score of a map/graph is calculated as the median of the distribution of module degree z-scores of all its (***g_global_(M))***. The vector (***g_bilateral_(M))*** is denoted by medians calculated separately over nodes on the ipsilateral and contralateral sides. Similarly, the median degree z-scores can be calculated on the resolution of microzones by pooling the nodes that lie within the zone ***g_zonewise_(M)***. The weighted module degree z-score is given by the following equation (Rubinov & Sporns, 2010):

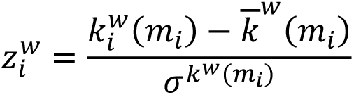 Where 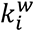 (*m_i_*) = weighted degree of node *i* with links between *i* and all nodes in *m_i_*; *m_i_*= module containing the node *i*; 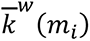 (*m_i_*) = mean of the within-module degree distribution; 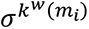 = standard deviation of the within-module degree distribution.
c. Participation coefficient (weighted) - measures how strongly does a node in a module connect to nodes of the other modules. Participation coefficient is close to 1 if the connections received by the node are uniformly distributed among all modules and close to 0 if it favors connections to the nodes within its own modules. A high participation coefficient for a graph indicates that patches in the map tend to be partially overlapping in their range or imperfectly separated, as islands in an archipelago linked by narrow “peers”. Participation coefficient is also a node centric measure, i.e., we get a value per node. The graph property on different resolutions (***g_global_(M), g_bilateral_(M) and g_zonewise_(M))*** are calculated as described for module degree z-score. The weighted participation coefficient is calculated as the equation (Rubinov & Sporns, 2010):

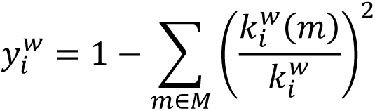 Where *M* = set of modules; 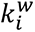(*m*) = weighted degree of links between node *i* and all nodes in module *m*; 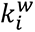 = weighted degree of links between node *i* and all nodes.
d. Local assortativity (weighted) - is a measure of the tendency of a node to connect to other nodes with the similar degree. A positive local assortativity coefficient indicates that nodes tend to link to other nodes with the similar degrees (e.g., if hubs tend to connect to hubs). Local assortativity is a node centric measure. Note that the node degree of a site within a map patch measures to how many other sites the response profile of the considered site is similar and that it is thus an indirect measure of the patch size scale. Thus, a map displaying both smaller and large patches well separated between them would result in a graph with a large assortativity. The graph property on different resolutions (***g_global_(M), g_bilateral_(M) and g_zonewise_(M))*** are calculated as described for module degree z-score. The weighted local assortativity is calculated with the following equation (Rubinov & Sporns, 2010, Leung et al, 2006):

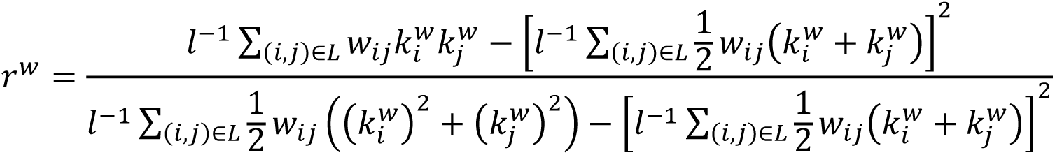 Where *l* total weight of all links in the network; *L*= set of links between pairs of nodes *i* and *j*; *w_ij_* = weight of the link between node *i* and *j*; 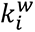 = weighted degree of node *i*; 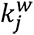 = weighted degree of node *j*.

#### Dimensionality reduction using t-SNE

t-SNE is a nonlinear dimensionality reduction technique that transforms vectors in a high dimensional metric space into vectors within a lower dimension metric space (usually 2 or 3 target dimensions for the sake of visualization) while ensuring that close points in the higher dimensional space are mapped to points which are still close in the lower dimensional space, whereas dissimilar points are mapped farther away. t-SNE projection were used to reduce the dimensionality for two kinds of data: (1) the zone wise distribution of synaptic weights (G_weights_zonewise_) and (2) lateralized values of graph properties (G_graph_bilateral_). The *sklearn.manifold.TSNE* function in python was used. Centroids of point clouds corresponding to maps obtained in similar locomotor adaptation conditions have been obtained by computing the median projected coordinates of all the maps in the considered point cloud.

#### Subtype Classification using random forest classifier

The graph features of different resolutions (***g_global_(M), g_bilateral_(M) and g_zonewise_(M))*** and zone wise synaptic weights (***gweights_zonewise(M)***) were used to train and test four random forest classifiers in order to classify the different locomotion contexts (CT, E/LC, E/LS, S/LTR). A random forest classifier uses multiple decision trees and combines their results to enhance the classification performance. The classifier used a maximum depth of 30 and 150 estimators. The data was divided into 100 stratified trials, with training:testing ratio of 80:20%. Proportion of correctly and misclassified samples (evaluated as generalization performance, i.e., applying the classifier on the testing subset not used for training) were quantified in a confusion matrix and the accuracy was calculated as a percentage of samples that were correctly classified. To compare the accuracy score with chance level scores, a shuffled version of the random forest was also trained and tested with the identical hyper parameters except the labels of the locomotion contexts in the data were shuffled. The *sklearn.ensemble.RandomForestClassifier* function from the sklearn package in python was used.

#### Generalized linear models

In order to check how well the behavioral features are predicted from the graph features, we trained and cross validated generalized linear models for 100 trials with training:testing split of 75:25%. The accuracy of the predictions on testing data (once again, generalization performance) was quantified by calculating r-values of fit between predicted and actual data for all 100 cross validation trials. To check the comparison with the chance level prediction, GLM was also trained and tested for the shuffled values of behavioral features in the training data. The r-value distributions of actual and shuffled GLMs are shown for comparison. The GLM equation posed the behavioral feature as a dependent variable and the graph-based features as independent variables. The contingency of graph features on the animal group (LC/LS or S-TR/L-TR) is encoded as a bias variable (±1, e.g., LC = 1, LS=-1) that is multiplied with the graph features:

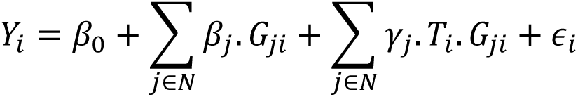

Where *i*= trial number; *Y*_*i*_ = behavioral feature for trial *i*; *β*_0_= common intercept; *N*= set of all graph-based features; *β*_*j*_= slope for graph feature *j*; *G_ji_* = value for graph property *j* in trial *i*; *γ*_*j*_= group specific slope for graph feature *j*; *T*_*i*_ = ±1, for the two groups (LC/LS & S/L-TR); ε_*i*_= error. *The statsmodels.api.GLM* routine from the python *statsmodels* package was used.

---

**Figure S1:**
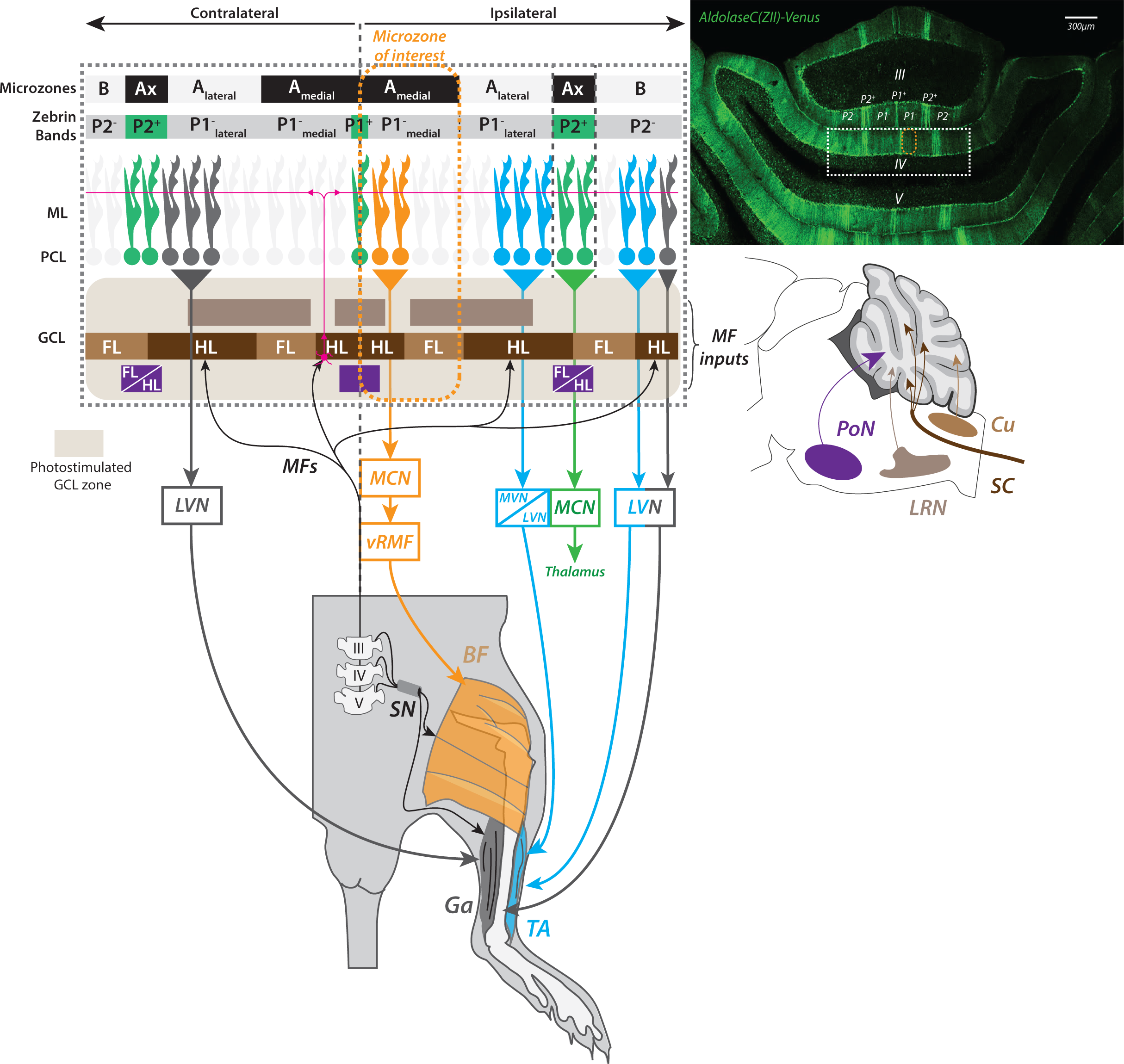
Input/output relationships in lobule II-IV of the anterior vermis. MF projections from precerebellar nuclei to the anterior microzones (A, AX, B) of the cerebellar cortex and microzonal organization based on Zebrin band patterning of PCs. Microzones target specific areas of cerebellar and vestibular nuclei. The medial cerebellar nuclei (MCN) project to the ventromedial reticular formation while the lateral vestibular nuclei can project directly to the lumbar spinal cord both controlling hindlimb muscles. Sensory information from hindlimb and forelimb muscles reaches the GCL in different microzones via MF originating in L3 to L5 lumbar segments and the cuneate nucleus, respectively. MFs relay on GCs (in red), that send paralle fibers through the transverse plane of the molecular layer contacting hundreds of PCs in several microzones. ***Adapted from Ji & Hawkes (1994); Ruigrok et al (2008); Valera et al (2016); Voogd et al (2016) & Biswas et al (2019)***. **III/IV/V**: lumbar segments, **BF**: biceps femoralis, **Cu**: Cuneate nucleus; **FL/HL**: forelimbs/hindlimbs, **Ga**: gastrocnemius, **GCs:** Granule cells; **MCN**: medial cerebellar nuclei, **MVN**: medial vestibular nuclei, **LVN**: lateral vestibular nuclei, **PoN**: pontine nuclei, **SC**: spinal cord, **SN**: sciatic nerve, **TA**: tibialis anterior, **VMRF**: ventromedial reticular formation.

**Figure S2:**
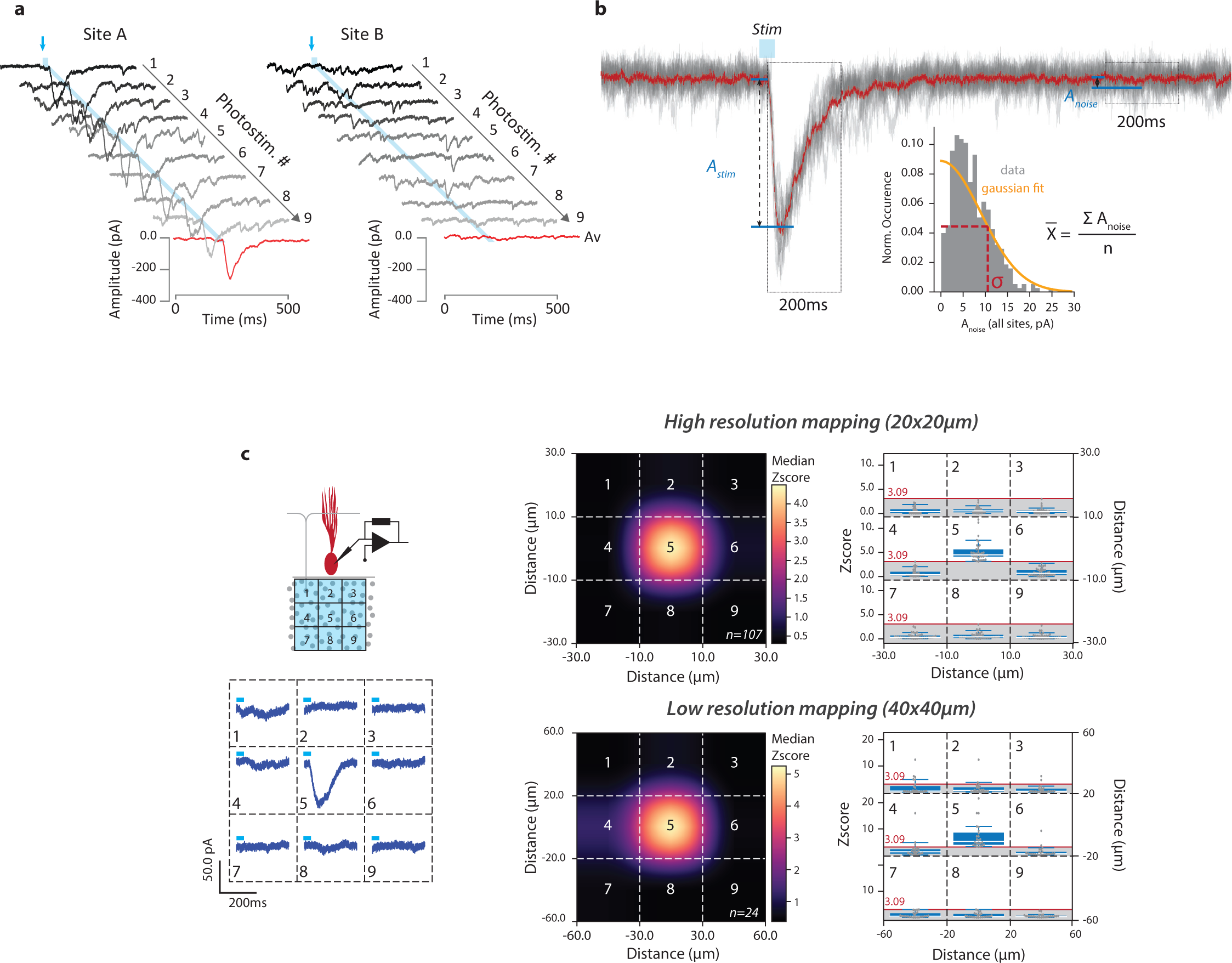
Glutamate uncaging methods. **(a)** Reliability of glutamate uncaging. Example traces of 9 consecutive photostimulation at 2 GCL sites (20*20µm, high resolution mapping, in shades of gray) and corresponding averaged EPSCs (in red). **(b)** Z Score calculation. In each GC site, we measured (1) the amplitude of the averaged response (A_stim_) in a 200ms time-win- dow following onset of light stimulation and (2) the amplitude of the averaged background noise (A_noise_ i.e. spontaneous GC activity). Distribution of A_noise_ values in a map (n=128 for low resolution and n=384 in high resolution. mappings) is (1) averaged (X) and (2) fitted with a gaussian kernel to extract standard deviation (σ). Z score of each GC site synaptic response (1 response per uncaging site) is then calculated as follows:

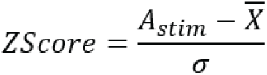 Significant photostimulation-evoked EPSCs in PCs (also called “active sites”) in the main text are determined via their individual z score (> 3.09 for active sites). **(c)** Spatial accuracy of GC-PC connectivity maps. In order to assess the spatial accuracy of RuBi-glutamate uncaging, we assessed whether we could identified isolated GC-PC sites in connectivity maps from CTRL (*high resolution, top right panel*) and P30-P40 (*low resolution, bottom right panel*) animals. 107 islands could be observed in high resolution (20*20µm) maps. Median of islands yielded following z scores: site #1: 0.35±0.49; site #2: 0.53±0.49; site#3: 0.36±0.34; site #4: 0.37±38; site #5 (center): 4.50±1.02; site #6: 0.66±0.65; site #7: 0.35±0.27; site #8: 0.41±0.34; site #9: 0.3±0.33. 24 islands could be identified in low resolution (40*40µm) maps. Median of islands yielded following z scores: site #1: 0.41±0.45; site #2: 0.69±0.84; site#3: 0.37±0.37; site #4: 1.29±1.27; site #5 (center): 5.25±2.45; site #6: 0.51±0.42; site #7: 0.5±0.47; site #8: 0.39±0.47; site #9: 0.41±0.32. Therefore, RuBi-glutamate does not diffuse outside individual illumination sites.

**Figure S3:**
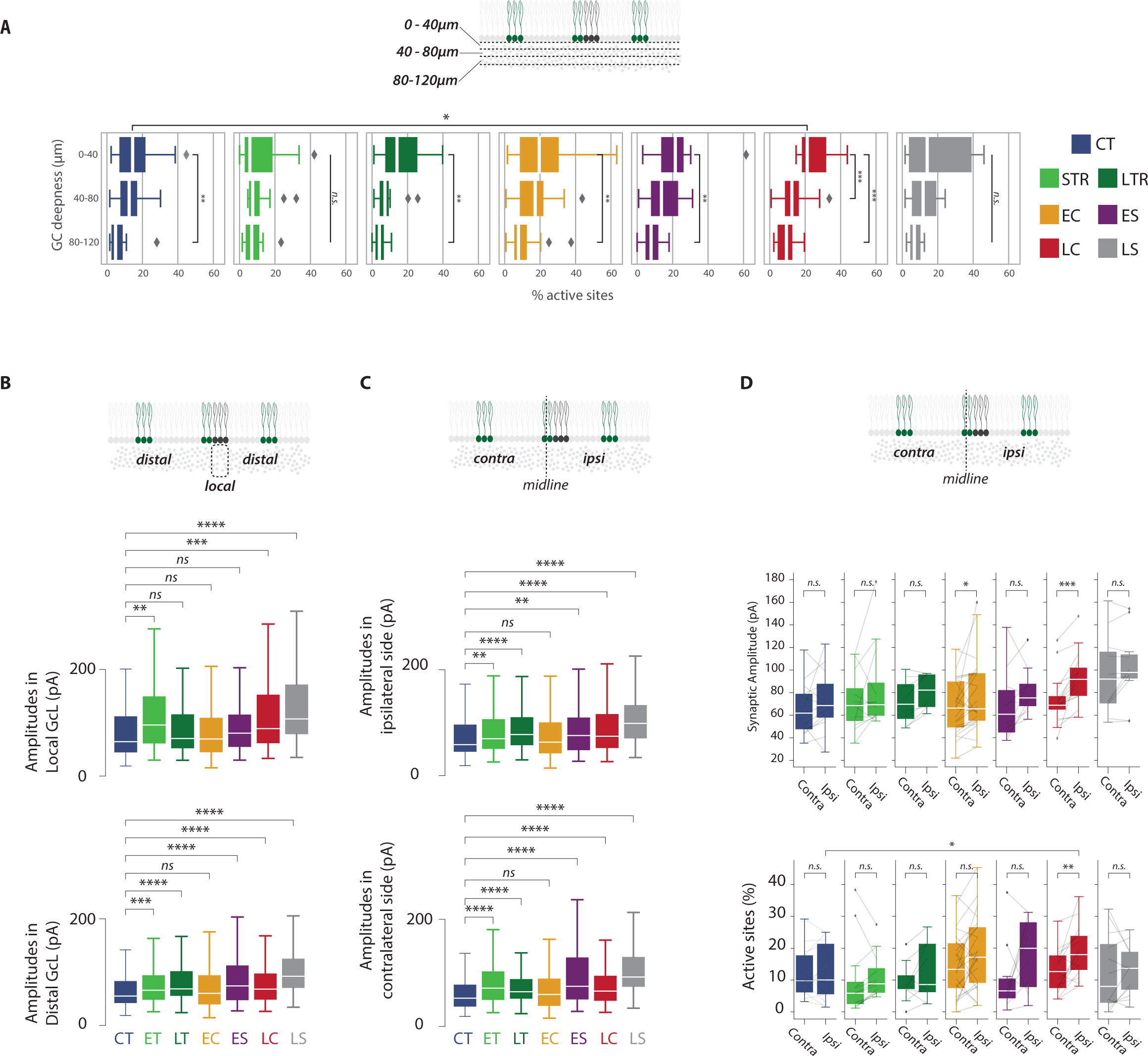
description of 2D maps. **(a)** Proportion of active sites in the height of the GC layer. GCL was subdivided in three zones from PcL to WM : 0 to 40µm, 40 to 80µm and 80 to 120µm layers. Proportion of active site was determined in each locomotor condition. Intra-group statistics were assessed with Kruskal Wallis test: CT, p=0.018; E-TR, p=0.60; L-TR,p=0.016; EC, p=0.0056; ES, p=0.031; LC, p=1.04e-05; LS, p=0.3. Post hoc Mann Whitney test: CT 0-40 vs 80-120, p=0.00934; L-TR 0-40 vs 80-120, p=0.0078; EC 0-40 vs 80-120 p=0.0029; ES 0-40 vs 80-120 p=0.011; LC 0-40 vs 40-80 p=5.10-4; LC 0-40 vs 80-120 p=1.10-5. Inter-group difference between CTRL and LC group (range 0-40µm) was assessed with Mann Whitney test (p=0.016). **(b)** Distribution of amplitude of active GC sites from local (top) and distal (down) GCL across locomotor condition. Local sites: pairwise comparisons were done with Mann Whitney U test after Kruskal Wallis test on all groups (p=1.10-6). Distal sites: pairwise comparisons were done with Mann Whitney U test after Kruskal Wallis test on all groups (p=7.10-18). **(c)** Same as in B for ipsilateral vs contralateral side. Ipsilateral: pairwise comparisons were done with Mann Whitney U test after Kruskal Wallis test on all groups (p=6.10-12). Contralateral: pairwise comparisons were done with Mann Whitney U test after Kruskal Wallis test on all groups (p=1.10-13). (D) Synaptic weights (top) and proportion of active sites (down) in the ipsi- and contralateral side of all synaptic mapsPairwise comparisons in each group were done with Wilcoxon Signed Rank test.

**Figure S4:**
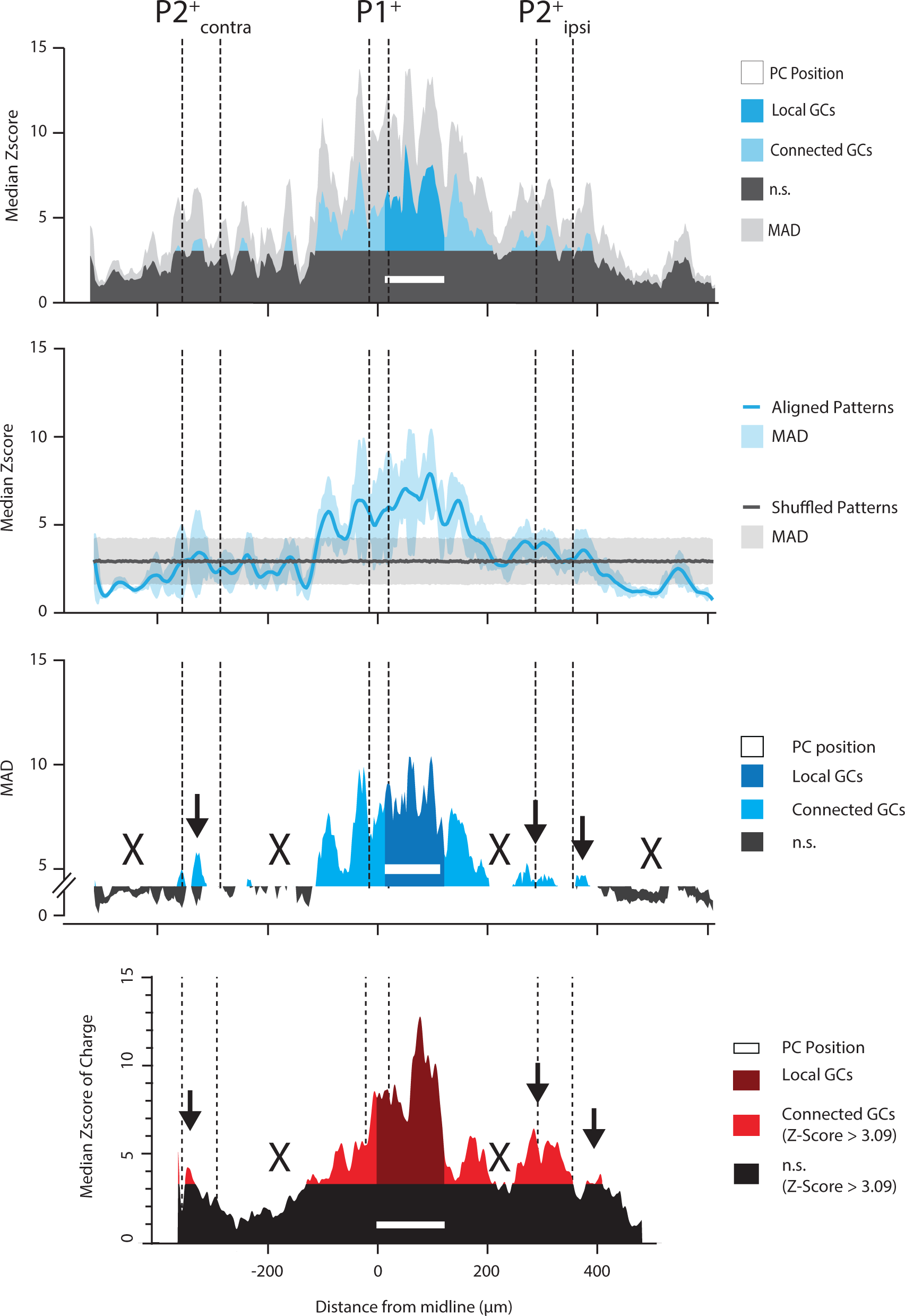
median 1D patterns of functional connectivity are conserved across animals. Individual synaptic 1D patterns used to build the median input pattern of medial PCs(first row) were bootstraped^81^ (average of 10000 unrestricted random samplings, second row). Patterns were either sorted by their coordinates along the mediolateral axis (blue curve) or shuffled (gray curve) removing positional information. Subtraction of Median Associated Deviations (MADs, third row) highlights connected GC sites above the randomized patterns (in blue) and silent areas (in black). Local GC input (in dark blue) and distant hotspots in both P2+ bands (black arrows) could be identified as well as silent areas in lateral P1- bands (black crosses), yielding a perfect match with previously described median input pattern (fourth row, adapted from^14^).

**Figure S5:**
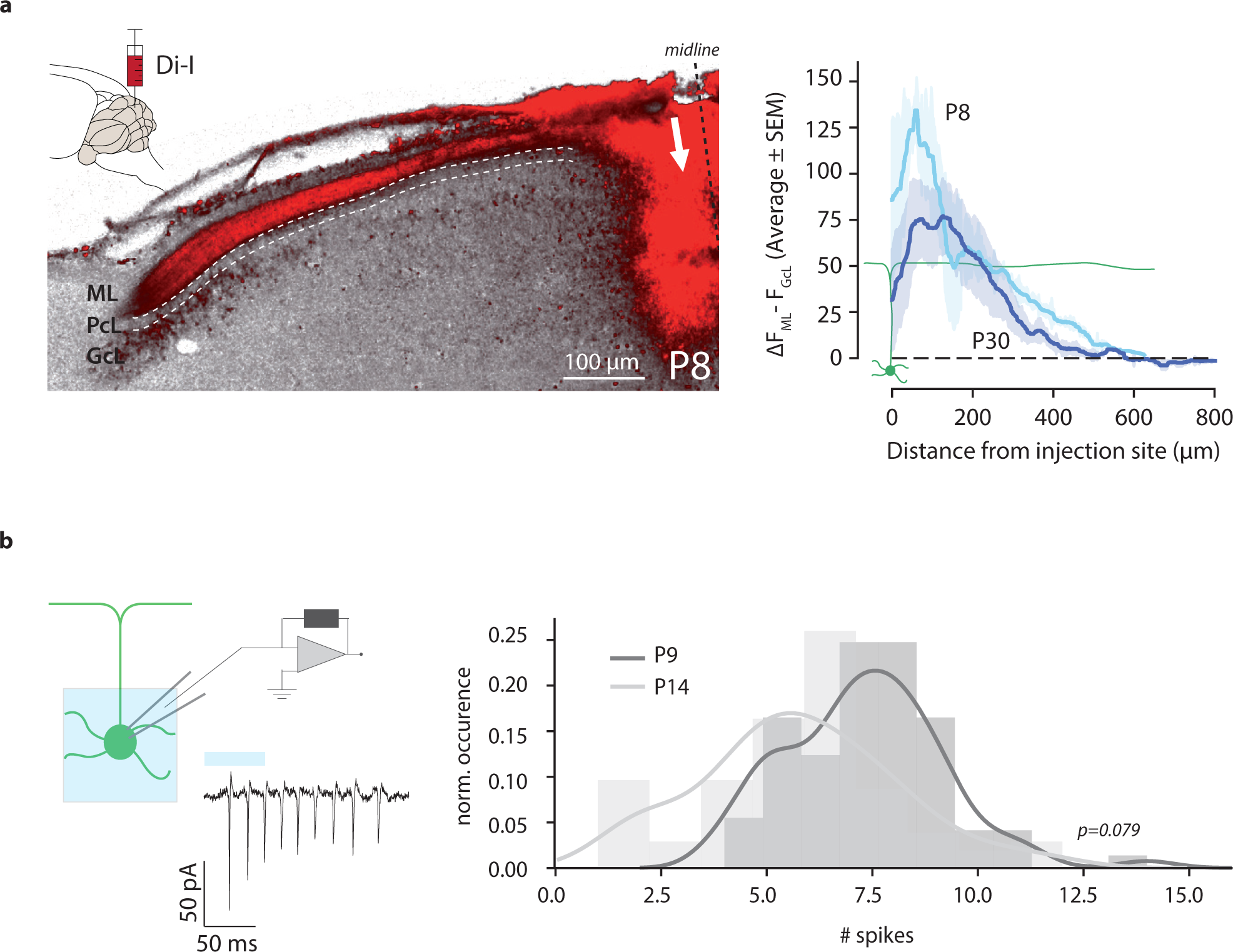
Parallel fibers length at PND8-30 and GC excitability. **(a)** Fluorescent Di-I was injected (Methods) at the midline of lobules III-V (left panel, injection site is shown with a white arrow). Fluorescence in ML and GCL were measured along the mediolateral axis from the injection site (right panel). **(b)** GCs excitability. Light-evoked action potentials in GCs were recorded via loose-cell attached method (left panel, Methods) in PND9 and PND14 mice. GCs (n=5 in both conditions) were randomly selected in the vermal GCL (lobules III, IV or V). Rubi-Glutamate uncaging triggered 7.3±0.8 action potentials in P9 pups and 5.7±1.3 action potentials in P14 mice (p=0.079, independent t-test). Reproducibility of GC firing patterns throughout consecutive photostimulation was previously assessed in^82^.

**Figure S6:**
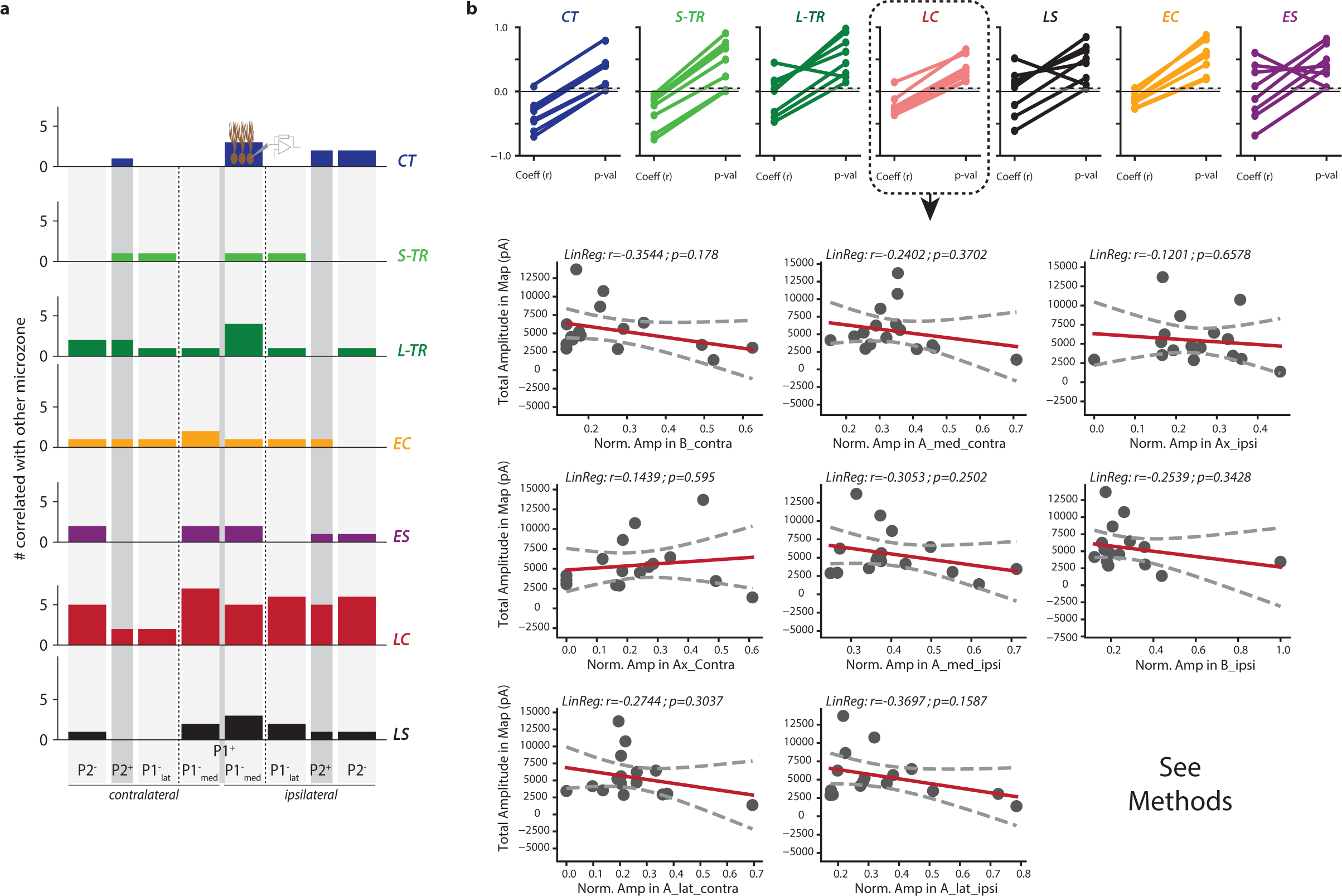
Zone wise Pearson correlation of synaptic weights. **(a)** Summary of zone wise Pearson synaptic correlation. For each microzone, we counted the amount of significant pairwise correlations with all the other microzones. Connectivity maps from the LC group showed a 5-fold increase in the number of correlations when compared to CTRL group (30 vs 6 correlations, with each microzone in LC group being correlated with 4.2 other zones on average). **(b)** Zone wise correlations are not the result of variability in slice GC overall excitability. We plotted the normalized amplitude of each microzone with the total synaptic weight recorded in the corresponding map. Amongst the 56 combinations, only 5 significant correlations (but negative, see top panel) were found in CT, STR, LS.

**Figure S7:**
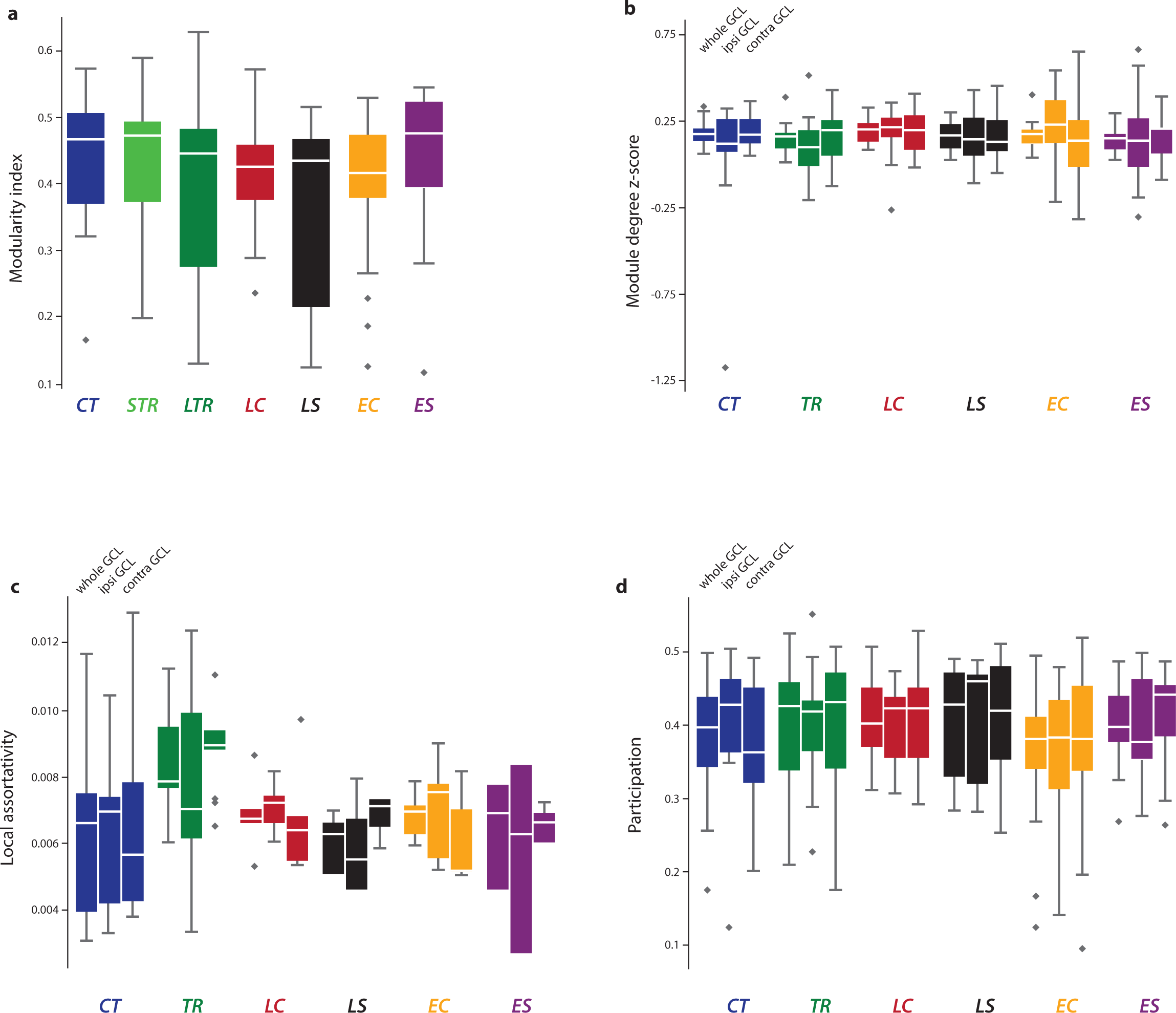
Graph parameters in all conditions. Distribution of global modularity index (**panel a**, i.e. whole map) and lateralized (i.e. from ipsi and contralateral side of the maps); module degree z score **(b)**, local assortativity **(c)** and participation **(d)** in each behavioral condition.

**Figure S8:**
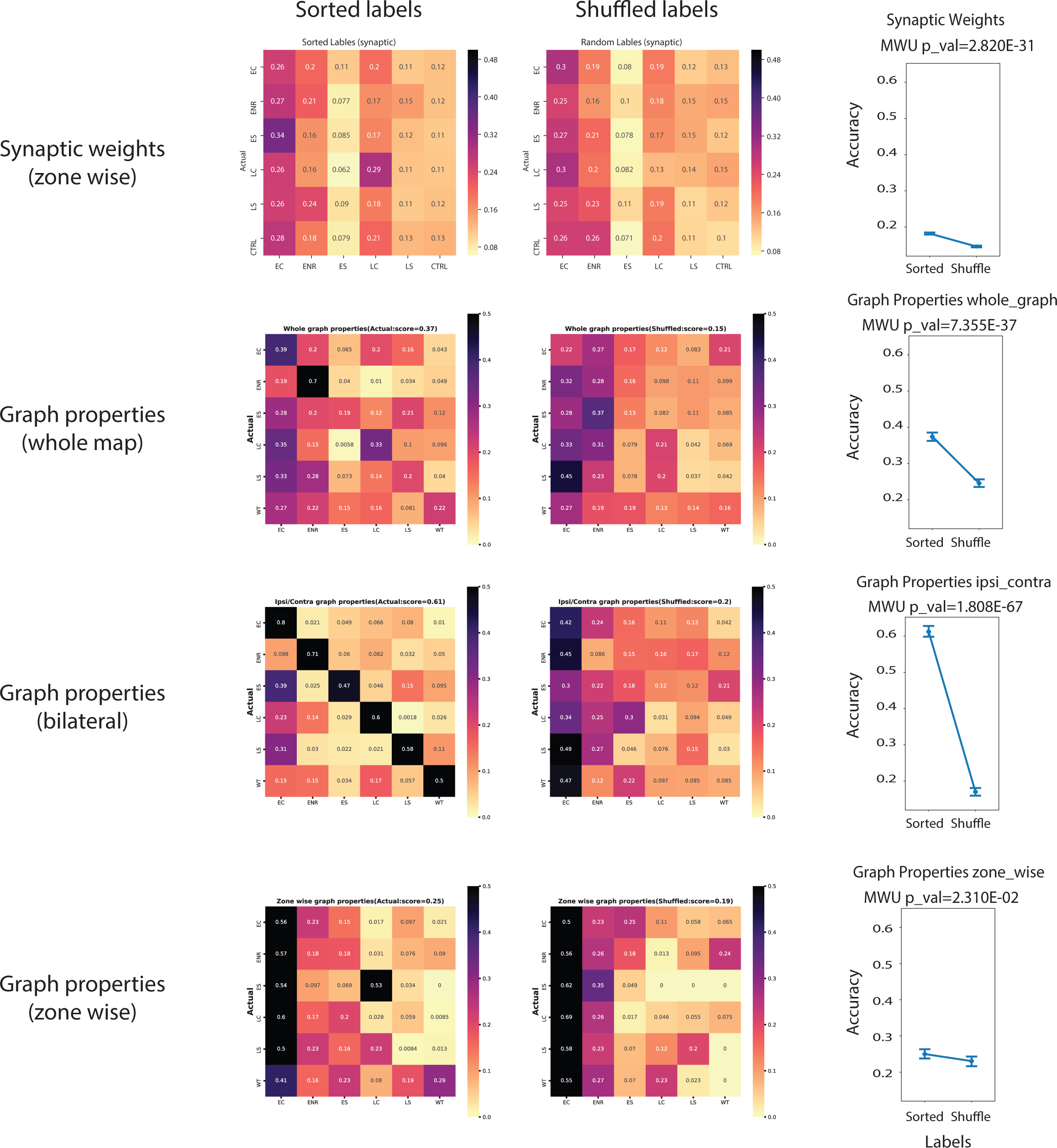
Random forest supervised classifications of synaptic maps. Confusion (i.e., correlation) matrices of Random Forest classification using input vector of zone wise synaptic weights (top), whole map graph properties (second row), bilateral graph properties (third row) and zone wise graph properties (fourth row). Matrices on the left were computed with correct labeling of the different vectors while matrices on the right result from shuffled input vectors (i.e., each vector was randomly labeled among CT/TR/ES/LS/LC/EC labels). Corresponding average accuracy is shown in the point plots on the right.

**Figure S9:**
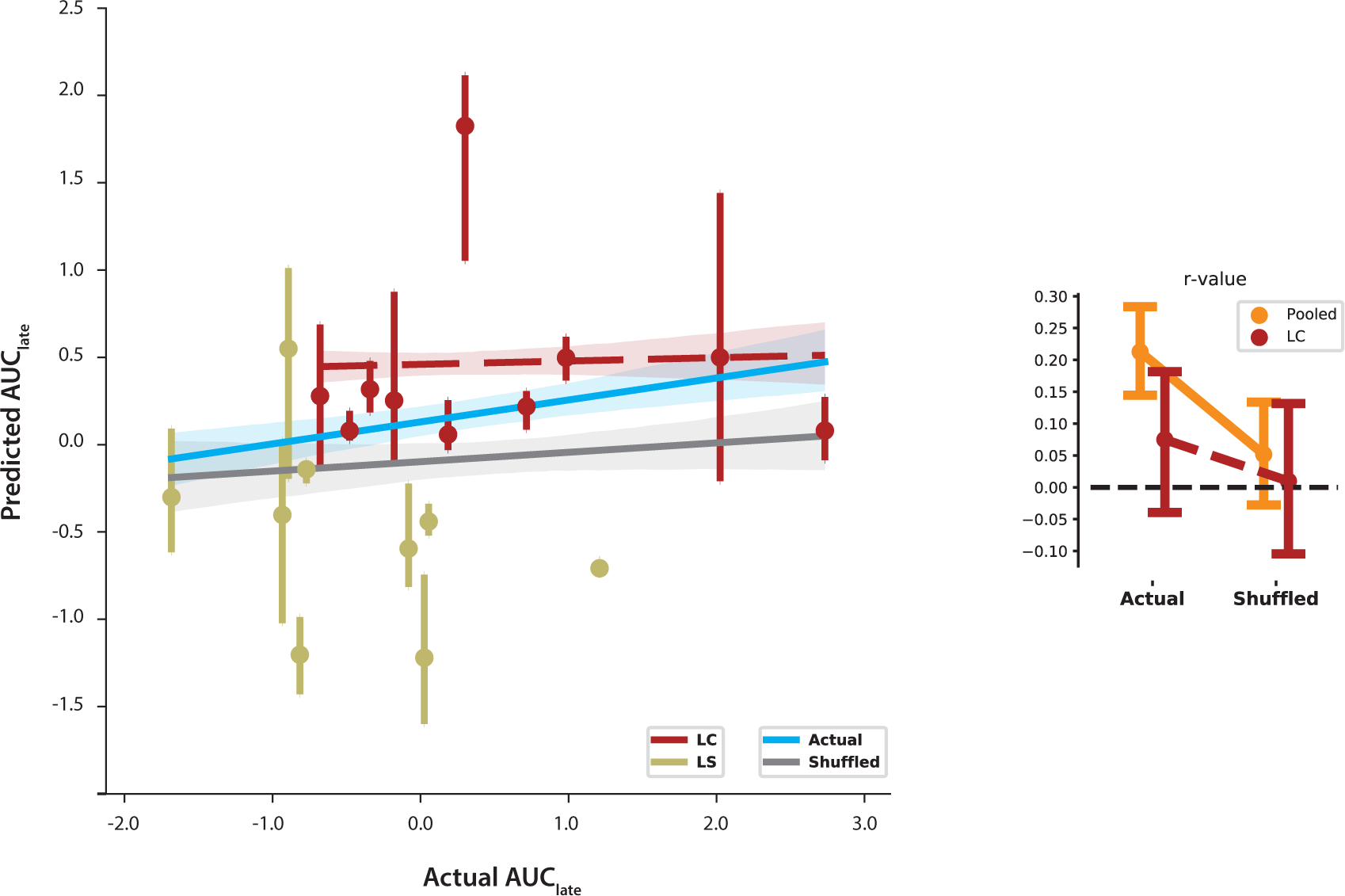
Example of failure for prediction using GLMs. Extended results of behavioral features prediction based on graph properties of synaptic maps using Generalized Linear Models (GLMs). When considering late post-surgery imbalance (AUC_late_) for prediction, GLMs cannot predict behavioral outcome based on graph properties. Average predictions are scattered on left panel, and regression coefficients (r value21s) are plotted in right panel. Linear models with actual data are shown in blue while results with shuffled data (i.e., chance level) are shown in gray. Analysis on LC group only is shown in dashed red.

**Figure S10:**
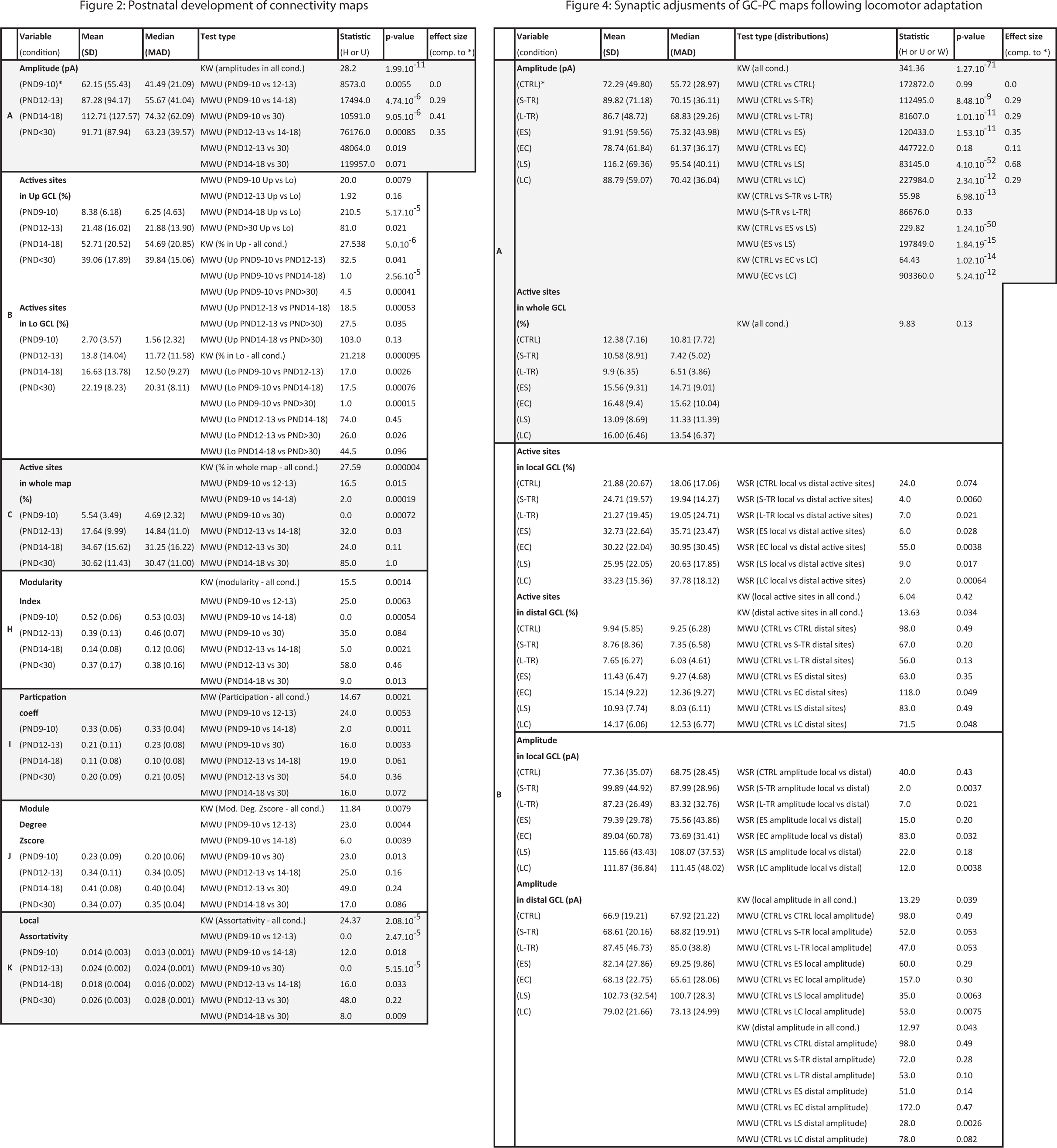
Statistic tables for. **Figure 2** **&** **Figure 4** Tables containing mean and median distributions as well as statistical metrics of different tests performed between distributions shown in Figure 2 (***left table***) and 4 (**right table**). Statistic H, U and W account for Kruskall-Wallis (KW), one-sided Mann-Whitney (corrected for ties) and two-sided Wilcoxon Signed-rank (WSR) tests. Effect size was calculate based on means (i.e., considers the standardized mean difference between two populations).

## References

1. Gao, Z. et al. A cortico-cerebellar loop for motor planning. Nature 563, 113–116 (2018).

2. Chabrol, F. P., Blot, A. & Mrsic-Flogel, T. D. Cerebellar Contribution to Preparatory Activity in Motor Neocortex. Neuron 103, 506–519 (2019).

3. Economo, M. N. et al. Distinct descending motor cortex pathways and their roles in movement. Nature 563, 79–84 (2018).

4. Li, N., Daie, K., Svoboda, K. & Druckmann, S. Robust neuronal dynamics in premotor cortex during motor planning. Nature 532, 459–464 (2016).

5. Sauerbrei, B. A. et al. Cortical pattern generation during dexterous movement is input- driven. Nature 577, 386–391 (2020).

6. Camon, J. et al. The Timing of Sensory-Guided Behavioral Response is Represented in the Mouse Primary Somatosensory Cortex. Cereb. Cortex 1–14 (2018) doi:10.1093/cercor/bhy169.

7. Shepherd, G. M. G., Pologruto, T. A. & Svoboda, K. Circuit analysis of experience- dependent plasticity in the developing rat barrel cortex. Neuron 38, 277–89 (2003).

8. Raymond, J. L. & Medina, J. F. Computational Principles of Supervised Learning in the Cerebellum. Annu Rev Neurosci 41, 233–253 (2018).

9. Petersen, C. C. H. The functional organization of the barrel cortex. Neuron vol. 56 339–355 (2007).

10. Collins, D. P. & Anastasiades, P. G. Cellular specificity of cortico-thalamic loops for motor planning. J. Neurosci. 39, 2577–2580 (2019).

11. Litwin-Kumar, A., Harris, K. D., Axel, R., Sompolinsky, H. & Abbott, L. F. Optimal Degrees of Synaptic Connectivity. Neuron 93, 1153–1164.e7 (2017).

12. Okun, M. et al. Diverse coupling of neurons to populations in sensory cortex. Nature 521, 511–515 (2015).

13. Portugues, R., Feierstein, C. E., Engert, F. & Orger, M. B. Whole-brain activity maps reveal stereotyped, distributed networks for visuomotor behavior. Neuron 81, 1328– 1343 (2014).

14. Valera, A. M. et al. Stereotyped spatial patterns of functional synaptic connectivity in the cerebellar cortex. Elife 5, e09862 (2016).

15. Coltz, J. D., Johnson, M. T. & Ebner, T. J. Cerebellar Purkinje cell simple spike discharge encodes movement velocity in primates during visuomotor arm tracking. J. Neurosci. 19, 1782–803 (1999).

16. Chen, S., Augustine, G. J. & Chadderton, P. The cerebellum linearly encodes whisker position during voluntary movement. Elife 5, 1–16 (2016).

17. Wolpert, D. M., Miall, R. C. & Kawato, M. Internal models in the cerebellum. Trends Cogn. Sci. 2, 338–47 (1998).

18. Bastian, A. J. Learning to predict the future: the cerebellum adapts feedforward movement control. Curr. Opin. Neurobiol. 16, 645–9 (2006).

19. Streng, M. L., Popa, L. S. & Ebner, T. J. Modulation of sensory prediction error in Purkinje cells during visual feedback manipulations. Nat. Commun. 9, 1099 (2018).

20. Ito, M. Control of mental activities by internal models in the cerebellum. Nat. Rev. Neurosci. 9, 304–13 (2008).

21. Shadmehr, R., Smith, M. A. & Krakauer, J. W. Error correction, sensory prediction, and adaptation in motor control. Annual Review of Neuroscience vol. 33 89–108 (2010).

22. Pasalar, S., Roitman, a V, Durfee, W. K. & Ebner, T. Force field effects on cerebellar Purkinje cell discharge with implications for internal models. Nat. Neurosci. 9, 1404–11 (2006).

23. Horn, K. M., Pong, M. & Gibson, A. R. Functional relations of cerebellar modules of the cat. J. Neurosci. 30, 9411–9423 (2010).

24. Powell, K., Mathy, A., Duguid, I. & Häusser, M. Synaptic representation of locomotion in single cerebellar granule cells. Elife 4, (2015).

25. Fujita, H., Kodama, T. & Du Lac, S. Modular output circuits of the fastigial nucleus for diverse motor and nonmotor functions of the cerebellar vermis. Elife 9, 1–91 (2020).

26. Morton, S. M. & Bastian, A. J. Mechanisms of cerebellar gait ataxia. Cerebellum 6, 79–86 (2007).

27. MacKinnon, C. D. Sensorimotor anatomy of gait, balance, and falls. in Handbook of Clinical Neurology vol. 159 3–26 (Elsevier B.V., 2018).

28. Pijpers, A., Winkelman, B. H. J., Bronsing, R. & Ruigrok, T. J. H. Selective impairment of the cerebellar C1 module involved in rat hind limb control reduces step-dependent modulation of cutaneous reflexes. J. Neurosci. 28, 2179–89 (2008).

29. Ruigrok, T. J. H., Pijpers, A., Goedknegt-Sabel, E. & Coulon, P. Multiple cerebellar zones are involved in the control of individual muscles: a retrograde transneuronal tracing study with rabies virus in the rat. Eur. J. Neurosci. 28, 181–200 (2008).

30. Apps, R. & Hawkes, R. Cerebellar cortical organization: a one-map hypothesis. Nat. Rev. Neurosci. 10, 670–81 (2009).

31. Michikawa, T. et al. Distributed sensory coding by cerebellar complex spikes. bioRxiv (2020) doi:10.1101/2020.09.18.301564.

32. Voogd, J. & Ruigrok, T. J. H. The organization of the corticonuclear and olivocerebellar climbing fiber projections to the rat cerebellar vermis: the congruence of projection zones and the zebrin pattern. J. Neurocytol. 33, 5–21 (2004).

33. Cerminara, N. L., Aoki, H., Loft, M., Sugihara, I. & Apps, R. Structural basis of cerebellar microcircuits in the rat. J. Neurosci. 33, 16427–42 (2013).

34. Oscarsson, O. Functional units of the cerebellum - sagittal zones and microzones. Trends Neurosci. 2, 143–145 (1979).

35. Garwicz, M., Ekerot, C.-F. & Jörntell, H. Organizational Principles of Cerebellar Neuronal Circuitry. News Physiol. Sci. An Int. J. Physiol. Prod. Jointly by Int. Union Physiol. Sci. Am. Physiol. Soc. 13, 26–32 (1998).

36. Gao, Z., van Beugen, B. J. & De Zeeuw, C. I. Distributed synergistic plasticity and cerebellar learning. Nat. Rev. Neurosci. 13, 619–35 (2012).

37. Ito, M. Bases and implications of learning in the cerebellum--adaptive control and internal model mechanism. Prog. Brain Res. 148, 95–109 (2005).

38. Jörntell, H. & Hansel, C. Synaptic memories upside down: bidirectional plasticity at cerebellar parallel fiber-Purkinje cell synapses. Neuron 52, 227–38 (2006).

39. Ly, R. et al. T-type channel blockade impairs long-term potentiation at the parallel fiber-Purkinje cell synapse and cerebellar learning. Proc. Natl. Acad. Sci. U. S. A. 110, 20302–7 (2013).

40. Shambes, G. M., Gibson, J. M. & Welker, W. Fractured somatotopy in granule cell tactile areas of rat cerebellar hemispheres revealed by micromapping. Brain. Behav. Evol. 15, 94–140 (1978).

41. Apps, R. et al. Cerebellar Modules and Their Role as Operational Cerebellar Processing Units. Cerebellum (2018) doi:10.1007/s12311-018-0952-3.

42. Shinoda, Y., Sugihara, I., Wu, H. S. & Sugiuchi, Y. The entire trajectory of single climbing and mossy fibers in the cerebellar nuclei and cortex. Prog. Brain Res. 124, 173–86 (2000).

43. Chabrol, F. P., Arenz, A., Wiechert, M. T., Margrie, T. W. & DiGregorio, D. A. Synaptic diversity enables temporal coding of coincident multisensory inputs in single neurons. Nat. Neurosci. 18, 718–27 (2015).

44. Huang, C.-C. et al. Convergence of pontine and proprioceptive streams onto multimodal cerebellar granule cells. Elife 2, e00400 (2013).

45. Fujita, H. et al. Detailed expression pattern of aldolase C (Aldoc) in the cerebellum, retina and other areas of the CNS studied in Aldoc-Venus knock-in mice. PLoS One 9, e86679 (2014).

46. Fino, E. et al. RuBi-Glutamate: Two-Photon and Visible-Light Photoactivation of Neurons and Dendritic spines. Front Neural Circuits 3, 2 (2009).

47. Ji, Z. & Hawkes, R. Topography of Purkinje cell compartments and mossy fiber terminal fields in lobules II and III of the rat cerebellar cortex: spinocerebellar and cuneocerebellar projections. Neuroscience 61, 935–54 (1994).

48. Gebre, S. A., Reeber, S. L. & Sillitoe, R. V. Parasagittal compartmentation of cerebellar mossy fibers as revealed by the patterned expression of vesicular glutamate transporters VGLUT1 and VGLUT2. Brain Struct. Funct. 217, 165–80 (2012).

49. Bollobás, B. Modern Graph Theory. vol. 184 (Springer New York,1998).

50. Rubinov, M. & Sporns, O. Weight-conserving characterization of complex functional brain networks. Neuroimage 56, 2068–2079 (2011).

51. Blondel, V. D., Guillaume, J. L., Lambiotte, R. & Lefebvre, E. Fast unfolding of communities in large networks. J. Stat. Mech. Theory Exp. 2008, P10008 (2008).

52. Wilson, H. A. et al. Low dose prenatal testosterone exposure decreases the corticosterone response to stress in adult male, but not female, mice. Brain Res. 1729, (2020).

53. Brust, V., Schindler, P. M. & Lewejohann, L. Lifetime development of behavioural phenotype in the house mouse (Mus musculus). Frontiers in Zoology vol. 12 S17 (2015).

54. Guan, W. et al. Eye opening differentially modulates inhibitory synaptic transmission in the developing visual cortex. Elife 6, (2017).

55. Bellardita, C. & Kiehn, O. Phenotypic Characterization of Speed-Associated Gait Changes in Mice Reveals Modular Organization of Locomotor Networks. Curr. Biol. 25, 1426–1436 (2015).

56. Breiman, L. Random forests. Mach. Learn. 45, 5–32 (2001).

57. Quy, P. N., Fujita, H., Sakamoto, Y., Na, J. & Sugihara, I. Projection patterns of single mossy fiber axons originating from the dorsal column nuclei mapped on the aldolase C compartments in the rat cerebellar cortex. J. Comp. Neurol. 519, 874–99 (2011).

58. Matsushita, M. Spinocerebellar projections from the lowest lumbar and sacral-caudal segments in the cat, as studied by anterograde transport of wheat germ agglutinin- horseradish peroxidase. J. Comp. Neurol. 274, 239–54 (1988).

59. Yaginuma, H. & Matsushita, M. Spinocerebellar projections from the thoracic cord in the cat, as studied by anterograde transport of wheat germ agglutinin-horseradish peroxidase. J. Comp. Neurol. 258, 1–27 (1987).

60. Wu, H. S., Sugihara, I. & Shinoda, Y. Projection patterns of single mossy fibers originating from the lateral reticular nucleus in the rat cerebellar cortex and nuclei. J. Comp. Neurol. 411, 97–118 (1999).

61. Biswas, M. S., Luo, Y., Sarpong, G. A. & Sugihara, I. Divergent projections of single pontocerebellar axons to multiple cerebellar lobules in the mouse. J. Comp. Neurol. 527, 1966–1985 (2019).

62. Murray, A. J., Croce, K., Belton, T., Akay, T. & Jessell, T. M. Balance Control Mediated by Vestibular Circuits Directing Limb Extension or Antagonist Muscle Co- activation. Cell Rep. 22, 1325–1338 (2018).

63. Voogd, J. Deiters’ Nucleus. Its Role in Cerebellar Ideogenesis. The Cerebellum (2015) doi:10.1007/s12311-015-0681-9.

64. Kiehn, O. Decoding the organization of spinal circuits that control locomotion. Nat Rev Neurosci 17, 224–238 (2016).

65. Jörntell, H. & Ekerot, C.-F. Reciprocal Bidirectional Plasticity of Parallel Fiber Receptive Fields in Cerebellar Purkinje Cells and Their Afferent Interneurons. Neuron 34, 797–806 (2002).

66. Shambes, G. M., Beermann, D. H. & Welker, W. Multiple tactile areas in cerebellar cortex: another patchy cutaneous projection to granule cell columns in rats. Brain Res. 157, 123–8 (1978).

67. Giovannucci, A. et al. Cerebellar granule cells acquire a widespread predictive feedback signal during motor learning. Nat. Neurosci. 20, 727–734 (2017).

68. Wagner, M. J., Kim, T. H., Savall, J., Schnitzer, M. J. & Luo, L. Cerebellar granule cells encode the expectation of reward. Nature 544, 96–100 (2017).

69. Knogler, L. D., Markov, D. A., Dragomir, E. I., Štih, V. & Portugues, R. Sensorimotor Representations in Cerebellar Granule Cells in Larval Zebrafish Are Dense, Spatially Organized, and Non-temporally Patterned. Curr. Biol. 27, 1288–1302 (2017).

70. Brown, I. E. & Bower, J. M. Congruence of mossy fiber and climbing fiber tactile projections in the lateral hemispheres of the rat cerebellum. J. Comp. Neurol. 429, 59– 70 (2001).

71. Bower, J. M. & Woolston, D. C. Congruence of spatial organization of tactile projections to granule cell and Purkinje cell layers of cerebellar hemispheres of the albino rat: vertical organization of cerebellar cortex. J. Neurophysiol. 49, 745–66 (1983).

72. Isope, P. & Barbour, B. Properties of Unitary Granule {Cell→Purkinje} Cell Synapses in Adult Rat Cerebellar Slices. J. Neurosci. 22, 9668–9678 (2002).

73. Sims, R. E. & Hartell, N. A. Differences in transmission properties and susceptibility to long-term depression reveal functional specialization of ascending axon and parallel fiber synapses to Purkinje cells. J. Neurosci. 25, 3246–57 (2005).

74. Walter, J. T., Dizon, M.-J. & Khodakhah, K. The functional equivalence of ascending and parallel fiber inputs in cerebellar computation. J. Neurosci. 29, 8462–73 (2009).

75. Lau, C. et al. Effects of perfluorooctanoic acid exposure during pregnancy in the mouse. Toxicol. Sci. 90, 510–518 (2006).

76. Wahlsten, D. A developmental time scale for postnatal changes in brain and behavior of B6D2F2 mice. Brain Res. 72, 251–264 (1974).

77. Schwartz, E. J. et al. NMDA receptors with incomplete Mg 2+ block enable low- frequency transmission through the cerebellar cortex. J. Neurosci. 32, 6878–6893 (2012).

78. Kano, M. & Hashimoto, K. Activity-dependent maturation of climbing fiber to purkinje cell synapses during postnatal cerebellar development. in Cerebellum vol. 11 449–450 (Cerebellum, 2012).

79. Hensch, T. K. Critical period plasticity in local cortical circuits. Nature Reviews Neuroscience vol. 6 877–888 (2005).

80. Porter, M. A., Onnela, J.-P. & Mucha, P. J. Communities in Networks.

81. Tononi, G., Edelman, G. M. & Sporns, O. Complexity and coherency: integrating information in the brain. Trends Cogn. Sci. 2, 474–84 (1998).

82. Hua, J., Xiong, Z., Lowey, J., Suh, E. & Dougherty, E. R. Optimal number of features as a function of sample size for various classification rules. Bioinformatics 21, 1509– 15 (2005).

83. Sneath, P. H. A. & Sokal, R. R. Numerical taxonomy. The principles and practice of numerical classification. (San Francisco, W.H. Freeman and Company., 1973).

84. Battaglia, D., Karagiannis, A., Gallopin, T., Gutch, H. W. & Cauli, B. Beyond the frontiers of neuronal types. Front. Neural Circuits 7, 13 (2013).

85. Carlsson, G. Topology and data. Bull. Am. Math. Soc. 46, 255–308 (2009).

86. Freund, J. et al. Emergence of Individuality in Genetically Identical Mice. Science (80-.). 340, 756–759 (2013).

87. Darmohray, D. M., Jacobs, J. R., Marques, H. G. & Carey, M. R. Spatial and Temporal Locomotor Learning in Mouse Cerebellum. Neuron 102, 217–231.e4 (2019).

88. Proville, R. D. et al. Cerebellum involvement in cortical sensorimotor circuits for the control of voluntary movements. Nat. Neurosci. 17, 1233–9 (2014).

89. Efron, B. & Tibshirani, R. An introduction to the Bootstrap. (1993).

90. Dorgans, K. et al. Short-term plasticity at cerebellar granule cell to molecular layer interneuron synapses expands information processing. Elife 8, 1–24 (2019).

91. Benbouzid, M. et al. Sciatic nerve cuffing in mice: A model of sustained neuropathic pain. Eur. J. Pain 12, 591–599 (2008).

92. Demšar, J. et al. Orange: Data Mining Toolbox in Python. J. Mach. Learn. Res. 14, 2349-2353 (2013).

